# CRISPR-pass: Gene rescue of nonsense mutations using adenine base editors

**DOI:** 10.1101/545723

**Authors:** Choongil Lee, Dong Hyun Jo, Gue-Ho Hwang, Jihyeon Yu, Jin Hyoung Kim, Se-eun Park, Jin-Soo Kim, Jeong Hun Kim, Sangsu Bae

**Affiliations:** Department of Chemistry, Seoul National University, Seoul 08826, South Korea; Center for Genome Engineering, Institute for Basic Science, Seoul 08826, South Korea; Fight against Angiogenesis-Related Blindness (FARB) Laboratory, Clinical Research Institute, Seoul National University Hospital, Seoul 03080, South Korea; Department of Chemistry, Hanyang University, Seoul 04763, South Korea; Research Institute for Convergence of Basic Sciences, Hanyang University, Seoul 04763, South Korea; Department of Biomedical Sciences, Seoul National University College of Medicine, Seoul 03080, South Korea; Department of Ophthalmology, Seoul National University College of Medicine, Seoul 03080, South Korea

**Keywords:** CRISPR-Cas9, base editing, nonsense mutation, stop codon read-through, premature termination codon

## Abstract

A nonsense mutation is a substitutive mutation in a DNA sequence that causes a premature termination during translation and produces stalled proteins resulting in dysfunction of a gene. Although it usually induces severe genetic disorders, there are no definite methods for inducing read-through of premature termination codons (PTCs). Here, we present a targeted tool for bypassing PTCs, named CRISPR-pass that uses CRISPR-mediated adenine base editors. CRISPR-pass, which should be applicable to 95.5% of clinically significant nonsense mutations in the ClinVar database, rescues protein synthesis in patient-derived fibroblasts, suggesting potential clinical utility.

## Introduction

Nonsense mutations, in which premature termination codons (PTCs) are formed by base pair substitution, truncate protein synthesis during translation. Such gene dysfunction is a source of severe pathological phenotypes in genetic diseases. Hence, compelling ribosomal read-through of the full coding sequence is a reasonable strategy for treating such genetic disorders. To address this issue, previous studies showed to induce the skipping of exons containing PTCs by using antisense oligonucleotides (AONs)^1, 2^. In other way, a few small-molecule drugs such as ataluren^3, 4^ and aminoglycosides^5^ (e.g., gentamicin) have been utilized to bypass nonsense mutations by introducing near-cognate tRNAs at the site of the PTC^6, 7^. However, those approaches act transiently and have nonspecific effects for the drugs. Alternatively, CRISPR-mediated homology-directed repair (HDR) can be used for gene correction, but is limited by low correction efficiency, especially in differentiated non-replicating cells from higher eukaryotes including humans^8, 9, 10^.

It was reported that CRISPR-mediated base editing technologies enable highly efficient direct conversion of DNA bases without producing double-strand breaks (DSBs). Cytidine deaminase-based base editors (BEs) produce C-to-T or G-to-A substitutions between the fourth and eighth bases in the non-binding strand of single-guide RNA (sgRNA) at protospacer DNA^11, 12^. On the other hand, A-to-G or T-to-C transitions in the same DNA positions can be achieved by adenine base editors (ABEs)^13^. In addition to the initial versions of BEs and ABEs, Koblan et al^14^ improved the base editing activities by expression optimization and ancestral reconstruction, which were named BEmax and ABEmax, respectively. Moreover, Hu et al^15^ and Nishimasu et al^16^ independently developed new Cas9 variants, named xCas9 and SpCas9-NG, that recognize 5’-NG-3’ with 5’-NAR-3’ sequences, expanding the targetable sites.

To date, a few groups reported to successfully correct target gene by restoring open reading frame in PTCs by using the ABEs such as a TAG-PTC of *EGFP* gene in rice^17^, both TAA-PTC of *Tyr* gene and TAG-PTC of *DMD* gene in mice^18^. However, although a few meaningful examples were shown, the systematic gene rescue for all possible cases are not demonstrated yet. It is expected that by targeting the coding strand with ABEs, the three possible PTCs, 5’-TAA-3’, 5’-TAG-3’, and 5’-TGA-3’, can be converted to 5’-TGG-3’, which will be translated to tryptophan (Trp). Alternatively, by targeting the noncoding strand, the three PTCs can be converted to 5’-CAA-3’ (translated to glutamine; Gln), 5’-CAG-3’ (Gln), or 5’-CGA-3’ (arginine; Arg), respectively (Fig. 1a). In this study, we established an ABE-mediated read-through method, named CRISPR-pass, to bypass PTCs by converting adenine to guanine or thymine to cytosine. We constructed all type of PTCs knock-in (KI) cell lines and then showed the read-through for all cases. Ultimately, we showed gene rescue at a patient-derived fibroblast containing PTC.

**Fig. 1.**
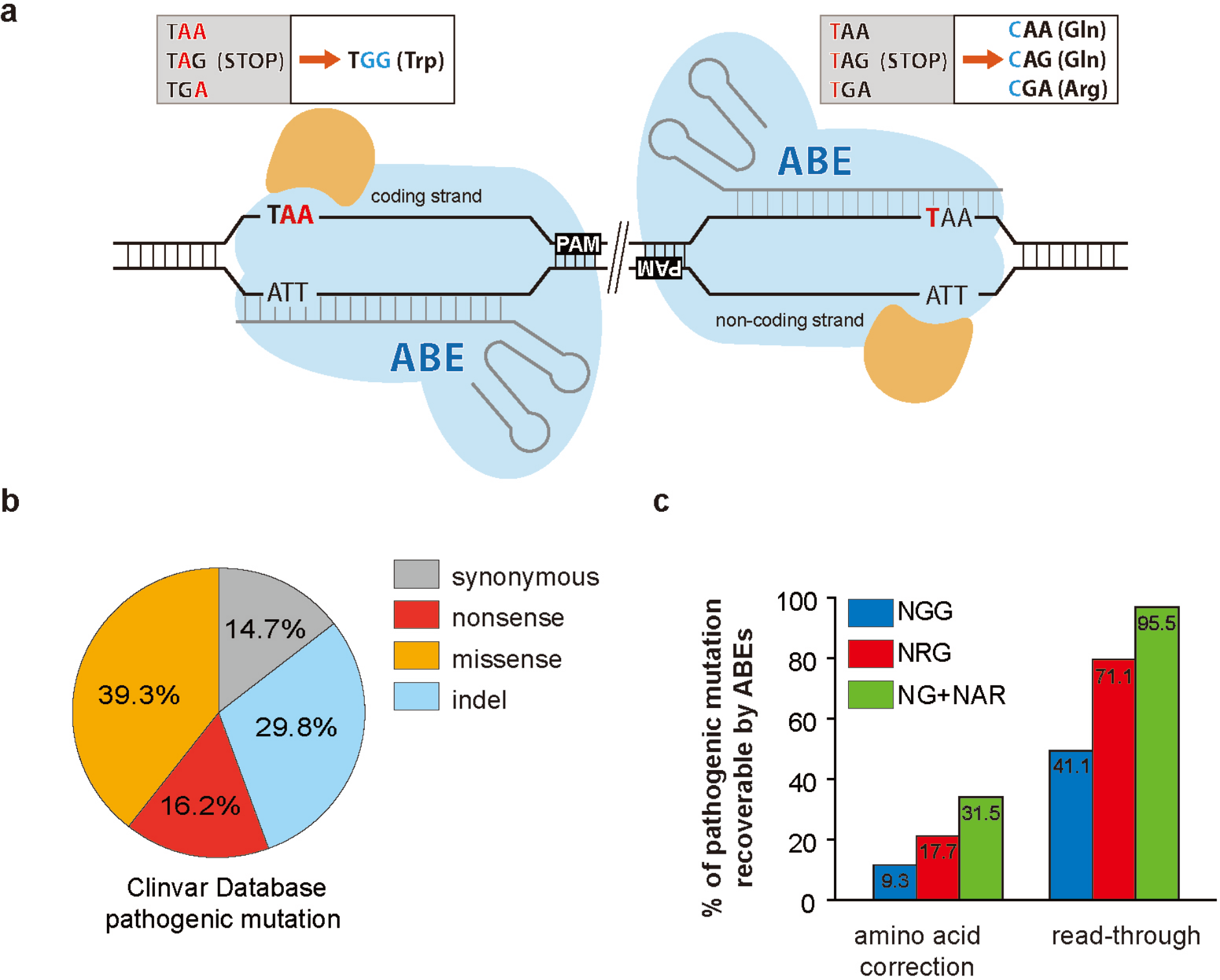
CRISPR-pass for restoring abbreviated gene expression. **(a)** Schematic of ABE-mediated CRISPR-pass. Targetable adenines are located in the coding or noncoding strand depend on the PAM’s orientation. All possible PTCs are shown in the upper boxes (coding strand targeting-TAA, TAG, TGA; noncoding strand targeting-TAA, TAG, TGA). The orange colored shapes represent adenosine deaminase. **(b)** The percentages of different types of mutations causing pathological phenotypes in the ClinVar database. **(c)** The percentages of PTCs that are targetable by CRISPR-pass with various PAM of variant ABEs and the recoverable rate of intact amino acids and bypassing alternative amino acids are depicted.

## Results

### *In silico* investigation of applicable targets for CRISPR-pass in the ClinVar database

We first inspected all targetable variations registered in the ClinVar database *in silico* to investigate how many genetic diseases with nonsense mutations could potentially be treated with CRISPR-pass. Of the 50,376 mutations causing pathological phenotypes in the database, nonsense mutations account for 16.2% (Fig. 1b); among these, 41.1 % are targetable by conventional ABEs that recognize a canonical protospacer adjacent motif (PAM), 5’-NGG-3’, and 95.5% are covered by a recently engineered ABE, xCas9 3.7-ABE 7.10 (called xABE here), which recognizes a non-canonical set of PAMs, 5’-NG-3’, and 5’-NAR-3’ (Fig. 1c). Only 31.5% of the nonsense mutations in the database can be exactly corrected to amino acids found in the non-mutant protein by xABEs, implying that the set of mutations that can be modified for read-through by CRISPR-pass is much larger than the set for which exact gene correction in DNA is possible (Fig. 1c and Supplementary Data 1).

### Construction of six knock-in HeLa cell lines carrying various types of PTCs in *EGFP* gene

To demonstrate the efficiency of CRISPR-pass in human cells, as a proof of concept, we tried to construct six knock-in HeLa cell lines, each carrying a different mutated version of the enhanced green fluorescent protein (*EGFP*) gene. We first prepared six DNA plasmids having different types of PTCs in the *EGFP* gene. The mutant *EGFP* genes as a set contain each type of PTC at two locations: the three PTCs in a position that can be converted by targeting the coding strand and the three PTCs in a position that can be converted by targeting the non-coding strand. The first position corresponds to a codon for lysine (Lys53) and the second to a codon for aspartate (Asp217); the encoded residues are located in connecting loop domains of EGFP (Fig. 2a and Supplementary Fig. 1). After preparing plasmids containing the six mutated *EGFP* genes, each plasmid was inserted into the genome in an endogenous safe-harbor region, the AAVS1 site^19^, using CRISPR-Cas9 via a non-homologous end-joining (NHEJ) pathway^20^ (Fig. 2b). The cell lines were named c-TAA, c-TAG, c-TGA, nc-TAA, nc-TAG, and nc-TGA, respectively.

**Fig. 2.**
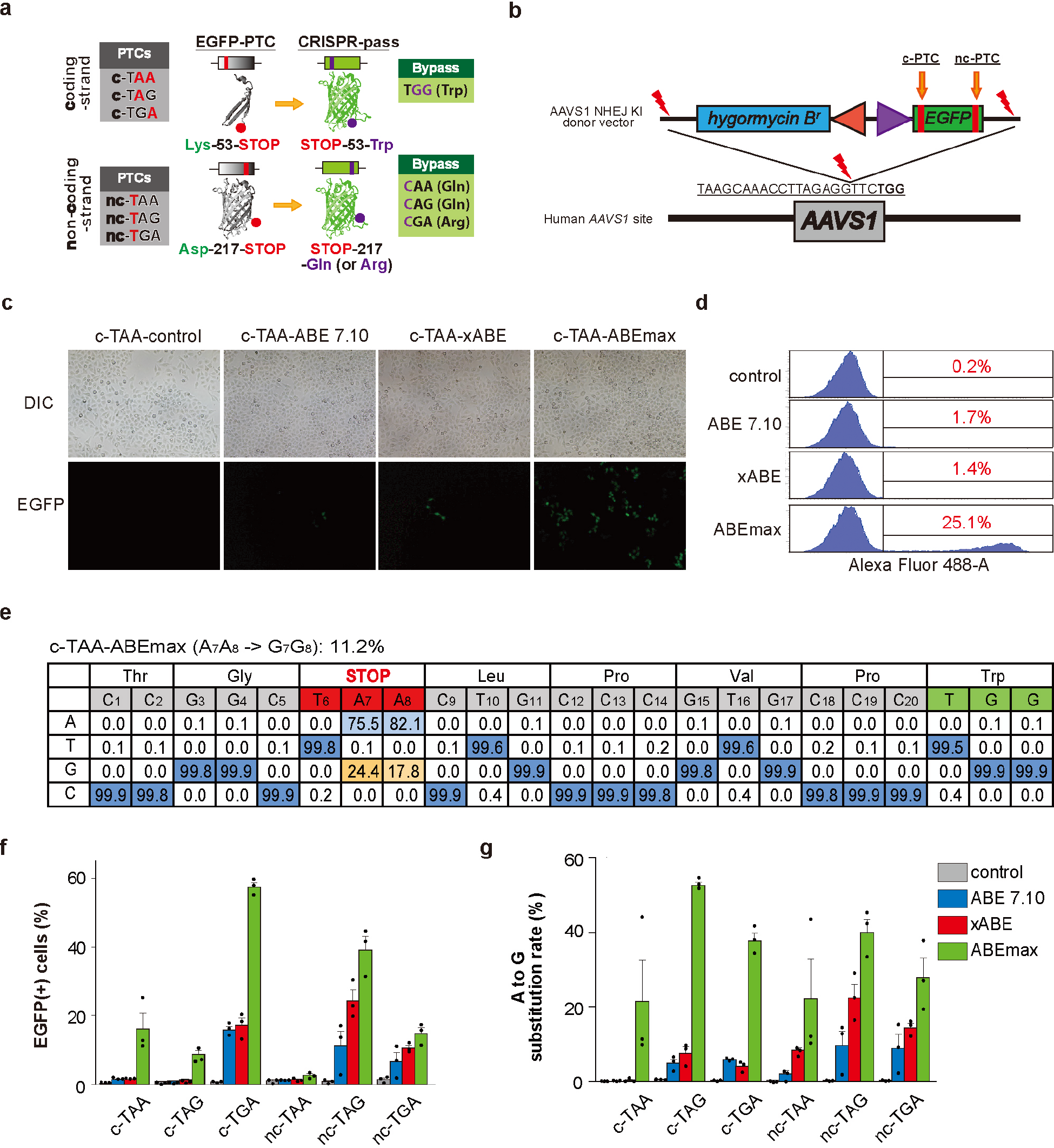
Restoring the function of *EGFP* gene expression in six KI HeLa cell lines carrying various types of PTCs. **(a)** Scheme for restoration of EGFP expression by CRISPR-pass. The first set of PTCs, which can be converted by targeting the coding strand, affect a residue that is located on a loop between the third and fourth beta strands; the second set of PTCs, which can be converted by targeting the noncoding strand, affect a residue that is located on a loop between the tenth and eleventh beta strands. c-PTC, coding strand PTC; nc-PTC, noncoding strand PTC. GFP structures was originated from the WIKIMEDIA COMMONS created by Zephyris. **(b)** Schematic of NHEJ-mediated KI of the EGFP-PTC constructs into the *AAVS1* site. Mutated EGFP KI cell lines were established for the three types of PTCs (TAA, TAG, and TGA). EGFP-PTC constructs were inserted into the *AAVS1* site by NHEJ-mediated KI methods. The hygromysin B resistant gene was also inserted for cell selection. **(c)** Fluorescence image of rescued EGFP expression in the c-TAA cell line after CRISPR-pass treatment. Three different versions of ABEs (ABE 7.10, xABE and ABEmax) were used for bypassing the PTCs in *EGFP* gene. **(d)** FACS analysis data after the different versions of ABEs (ABE 7.10, xABE and ABEmax) were treated in the c-TAA cell line. **(e)** Targeted deep sequencing data showing the percentages of each of the 4 nucleotides at each position in the target DNA sequences as a substitution table, which was obtained from the c-TAA cell line after the ABEmax treatment. Bar graphs showing recovered EGFP expression levels as determined by FACS analysis **(f)** and showing A-to-G substitution rates at PTC sites as determined by targeted deep sequencing **(g)** for each EGFP-PTC KI cell line, after treatment with ABEs (ABE 7.10, xABE, or ABEmax). Each dot represents the three independent experiments. Error bars represent s.e.m.

### CRISPR-pass rescues the function of *EGFP* gene in six knock-in HeLa cell lines

To test whether ABE treatment would allow bypass of these nonsense mutations, we transfected plasmids expressing sgRNAs designed to target both locations harboring PTCs, together with ABE-encoding plasmids, into the prepared HeLa cell lines. After ABE treatment by lipofection, we found that functional EGFP was expressed, as seen by green fluorescence, in all cell lines. For example, in the case of c-TAA cells, the function of EGFP would be rescued when two adenines are changed to guanines simultaneously for bypassing the PTC. As shown in Figure 2c, the functional EGFPs were observed after various ABEs (ABE 7.10, xABE, and ABEmax) were treated. We quantified the ratios of rescued to mutated EGFPs by fluorescence-activated cell sorting (FACS) analysis (Fig. 2d). We also confirmed the A to G conversion at target DNA region by targeted deep sequencing in bulk cell populations; the conversion rate of two adenines (A_7_A_8_) to two guanines (G_7_G_8_) was 11.2% here (Fig. 2e).

Similar to the c-TAA cells, we repeatedly carried out CRISPR-pass for all other types of KI cell lines (Supplementary Fig. 2). The quantitative ratios of the functional EGFP expression were also measured by FACS and targeted deep sequencing. As a result, the FACS analysis demonstrated that nonsense mutations were bypassed in 0.7-17.8% of cells (Fig. 2f, Supplementary Fig. 3 and Supplementary Table 1). And targeted deep sequencing analysis confirmed that 0.2-15.2% of the cells showed A-to-G conversions at target regions with a comparable tendency (Fig. 2g and Supplementary Table 2). It is noteworthy to mention that ABEmax was the most effective one in all cases, resulting in up to 59.6% rescue of the mutant *EGFP* gene, compared to the ABE7.10 and xABE.

**Fig. 3.**
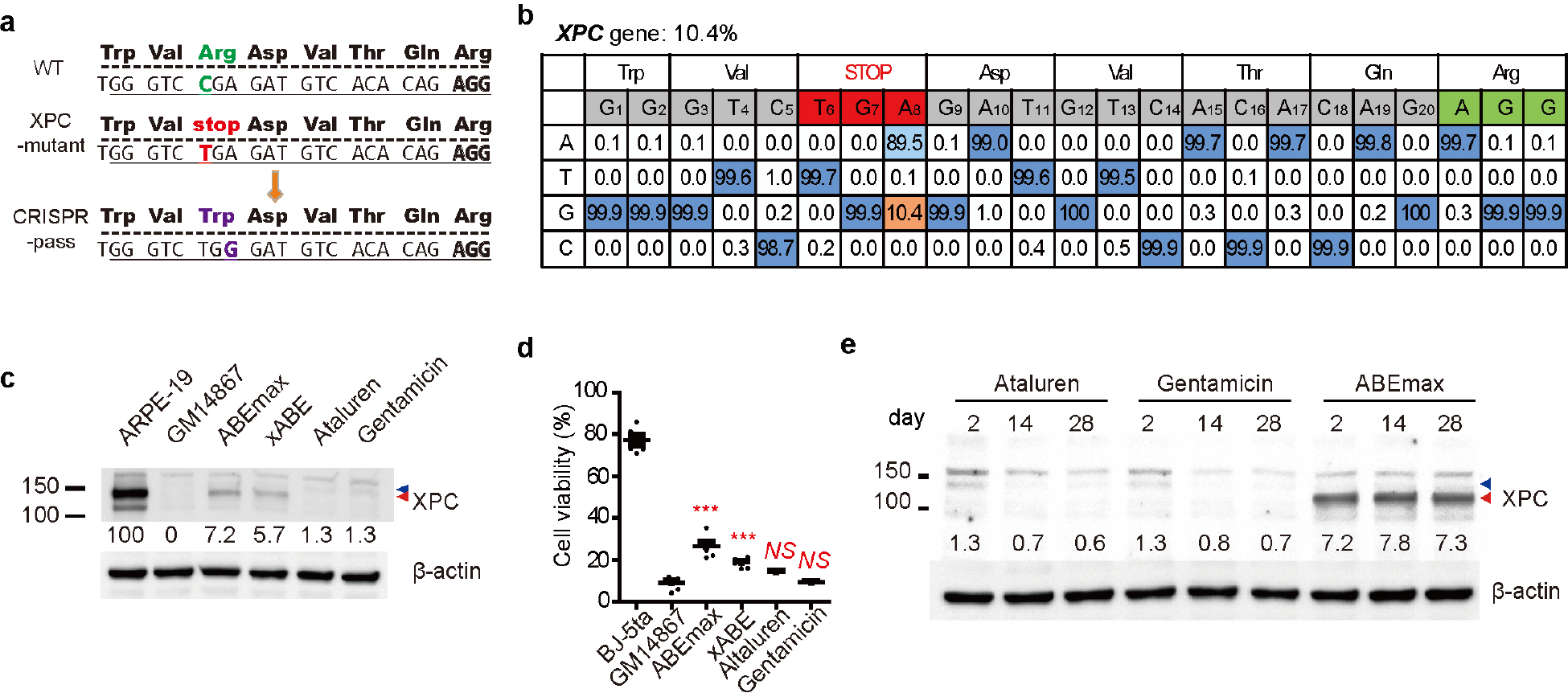
Restoring abbreviated *XPC* gene expression in patient-derived fibroblasts. **(a)** Scheme for ABE-induced read-through of an XPC-associated PTC. **(b)** Targeted deep sequencing data showing the A-to-G substitution rate induced by ABEmax treatment at the PTC site in the *XPC* gene. **(c)** Expression level of the XPC protein in *XPC* mutant cells rescued by treatment with ABEs (ABEmax or xABE), compared with the expression level in untreated cells and cells treated with ataluren or gentamicin for 48 h. **(d)** Cell viability of WT skin fibroblasts (BJ-5ta), *XPC* mutant cells (GM14867), and *XPC* mutant cells treated with ABEs (ABEmax or xABE), ataluren, or gentamicin at 3 days after exposure to 254 nm ultraviolet radiation at a dose of 25 J/m^2^. *P*-values were calculated by one-way ANOVA with post-hoc Bonferroni’s multiple comparison tests (*n* = 6). *P-*values indicators from a comparison with GM14867 cell viability are shown above each treatment group. *NS*, not significant (*P* > 0.05); *, *P* < 0.05; ***, *P* < 0.001. **(e)** Prolonged expression of the XPC protein after CRISPR-pass treatment. Significant and stable XPC protein expression was observed until at least 4 weeks after ABEmax treatment. However, XPC protein expression declined after removal of ataluren and gentamycin. Proteins were also prepared from ABEmax-treated XPC mutant cells at 2 and 4 wk (subculturing twice per week) for comparison. Blue and red arrowheads indicate the positions of XPC protein.

### CRISPR-pass rescues the function of *XPC* gene in patient-derived fibroblasts

We next applied CRISPR-pass to the rescue of a nonsense mutation in fibroblasts (GM14867) derived from a patient with xeroderma pigmentosum, complementation group C (XPC). XPC, which affects the skin, is a genetic disorder caused by nonsense mutations in the *XPC* gene. The XPC protein is an initiator of global nucleotide excision repair^21^. Thus, XPC-deficient cells accumulate DNA damage when they are exposed to chemical or physical stimuli including ultraviolet irradiation^22^. GM14867 cells have a homozygous C>T nonsense mutation at nucleotide 1840 in the *XPC* gene, which creates a 5’-TGA-3’ stop codon that replaces a codon for arginine (Arg579) (1840C>T, Arg-579-UGA stop codon) (Fig. 3a). After treating GM14867 cells with ABE7.10-encoding plasmid and sgRNA-encoding plasmid by electroporation, the adenine base in the 5’-TGA-3’ stop codon was converted to guanine to create 5’-TGG-3’ at a rate of 3.4%, as measured by targeted deep sequencing (Supplementary Fig. 4), indicating partial rescue of the *XPC* gene. Similar to the previous experiments, xABE and ABEmax resulted in higher base editing rates more than 10% (Supplementary Fig. 4 and Fig. 3b), respectively. Western blot analyses demonstrated that both xABE and ABEmax treatment led to recovery of expression of the full-length XPC protein, with a molecular weight similar to that in wild-type (WT) cells (ARPE-19), at considerably higher levels than induced by ataluren or gentamicin (Fig. 3c).

Next, to determine the functional activity of the recovered XPC protein, we evaluated the viability of GM14867 cells at 72 hours after exposure to 254 nm ultraviolet light. To our surprise, both xABE- and ABEmax-treated GM14867 cells had significantly regained resistance to ultraviolet irradiation-induced DNA damage, causing an increase in cellular viability (Student’s *t*-test, P<0.001, Fig. 3d). More importantly, ABEmax-treated GM14867 cells sustained such XPC protein expression for at least 4 weeks, whereas the cells treated with ataluren or gentamicin gradually lost XPC protein expression (Fig. 3e and Supplementary Fig. 5), implying that CRISPR-pass inducing persistent expression for the nonsense mediated disease therapies.

Finally, to examine the off-target effects of CRISPR-pass in GM14867 fibroblasts, we searched for potential off-target sites using Cas-OFFinder^23^ and carried out targeted deep sequencing for 12 candidate target sites (Supplementary Fig. 6 and Supplementary Table 3). As a result, we found no noticeable off-target sites likewise to the previous ABE-based gene editing studies^24, 25, 26^, suggesting potential clinical utility.

## Discussion

Previously, Kuscu et al^27^ and Billon et al^28^ demonstrated gene silencing method, named CRISPR-STOP and iSTOP, respectively, through BE-induced nonsense mutations. In this study, we analogously demonstrated that CRISPR-pass is a straightforward method for inducing read-through of PTCs by ABEs, covering most (95%) nonsense mutations in the ClinVar database that cause pathological phenotypes. We first demonstrated the CRISPR-pass activities in six types of EGPF-PTCs-KI human cells, as a proof of concept. In these experiments, CRISPR-pass efficiently rescued functional EGFP expression by bypassing all PTCs. Then, we successfully confirmed the activity of CRISPR-pass in a patient derived fibroblast, GM14867, which contains a nonsense mutation at *XPC* gene.

Until now, researchers have tried to correct the PTC in *XPC* coding gene by various approaches. One suggested a viral delivery method of intact *XPC* coding plasmids^29^, but it has potential problems such as a random integration of the transgene in viral delivery^30^ and overexpression effects of the exogenous *XPC* gene. Alternatively, another approach to correct endogenous *XPC* gene was demonstrated by using meganucleases and TALENs^22^. In this study, the authors tried to correct the *XPC* gene via a homology directed repair (HDR) pathway after producing double strand breaks (DSBs) of DNA, which might induce DSBs-mediated cell apoptosis^31^, whereas the CRISPR-pass does not generate DSBs of DNA. Furthermore, we showed that the A-to-G conversions at a rate of about 10% can rescue the expression of functional XPC protein (Fig. 3b-3d) without detectable off-target effects, strongly indicating that the CRISPR-pass is a relevant approach for rescuing the nonsense associated diseases with higher editing efficiencies than using HDR^32^ and without the loss of large portion of protein via the exon removal^33^ or skipping strategies^34^. More importantly, CRISPR-pass induced prolonged XPC protein expression unlike ataluren and gentamycin that are known as current nonsense mutation disease therapy^3, 5^.

Recently, it is reported that DNA cleavages at on-target site frequently cause undesired large deletions or complex genomic rearrangements^35^. In this aspect, CRISPR-pass has important safety advantages relative to approaches that do rely on DNA cleavage. Furthermore, recent off-target profiling experiments on ABEs supported the high specificity of ABEs^24, 25, 26^, increasing the potential clinical utility of it. These characteristics suggest that CRISPR-pass might be useful for gene rescue in a clinical setting, as an alternative to existing drugs.

## Methods

### General methods and cloning

All kinds of ABEs were purchased from addgene (pCMV-ABE 7.10, #102919; xCas9(3.7)-ABE(7.10), #108382; pCMV-ABEmax, #112095). The pXY-ECFP-AAVS1-NHEJ-KI donor vector (Plasmid^36,37^ was provided by prof. Woo. J. S. at Korea University in Republic of Korea, and it was modified by Jihyeon Yu who is one of the authors in this study) was digested with SacI and BsrGI, and Gibson assembly was then used to generate plasmids containing mutated versions of *EGFP*. The linearized vector was incubated with amplified *EGFP* DNA sequences, respectively containing each PTC at the appropriate location, with 20 nucleotides of overhanging homologous sequence at either end (5’- GGTCTATATAAGCAGAGCTC-3’ and 5’- TGTGCGGCTCACTTGTACAG-3’), in a solution containing 2X Gibson master mixture at 50°C for 1hour^38^. After then, pXY-EGFP-AAVS1-NHEJ-KI vector digested again with SacI. The linearized vector was incubated with additional amplified DNA sequences (ggtctatataagcagagctctcgtcgacgagctcgtttagtgaaccgtcagatcgtttaaacaagttggtcgtgaggcactgggcaggtaagtatcaaggttacaagacaggtttaaggagaccaatagaaactgggcttgtcgagacagagaagactcttgcgtttctgataggcacctattggtcttactgacatccactttgcctttctctccacaggtgtccagggtaccgagctcgccgccatggtgag) and 2x Gibson master mixture in 50°C. Each sequence of oligos encoding sgRNA was purchase from Macrogen Inc (South Korea). Oligos were heated and cooled down by a thermocycler for complementary annealing. Double strand oliges were ligated into linearized pRG2 plasmid linearized by BsaI restriction enzyme (addgene, #104274). List of oligomers for sgRNAs and Primer sequences for cloning are in Supplementary Table 4 and Supplementary Table 5.

### ClinVar database analysis

Bioinformatic analysis of the ClinVar database of human disease-associated mutations was conducted using Python. The ClinVar database (Common_and_clinical_20170905) was used for this analysis. The Python script used to analyze mutation patterns in human diseases and to identify mutations that could be CRISPR-pass targets can be accessed at website (https://github.com/Gue-ho/CRISPR-pass). Briefly, the steps of the analysis were as follows:

1-1) Among entries in the ClinVar database, we identified mutation patterns in the following categories: indels (insertions or deletions), silent mutations, nonsense mutations, and missense mutations. 1-2) For precise analysis, entries in each mutation pattern category were subdivided depending on their nucleotide sequence using information about the surrounding genomic sequence and coding sequence (CDS). CDSs were extracted from the SNP database at NCBI. If no CDS was found in NCBI than data were taken from GRCh38 and hg19.

2-1) Among the sorted entries from the ClinVar database, the number of nonsense mutations that were potential CRISPR-pass targets were counted. The targetable Cas9 sites were grouped by their associated PAM sequences, such as GG, AG, GA, GC, GT, GAN, and AA. 2-2) Each Cas9-targetable site was filtered depending on the ABE target range (position 4 to 8 in the protospacer from the end distal to the PAM) it contained. 2-3) To prevent counting sequences more than once, in the case of SpCas9, a sequence was counted when at least either GG or AG was possible; the number was counted as a targetable PAM for xABE when at least one of the PAM was possible.

### Cell culture and transfection

HeLa (ATCC, CCL-2) cells were grown in Dulbecco’s Modified Eagle Medium (DMEM) with 10% fetal bovine serum (FBS) and a penicillin/streptomycin mix (100 units/mL and 100 mg/mL, respectively). 2.5 × 10^5^ HeLa cells were transfected with each ABE (ABE, xABE, or ABEmax)-encoding plasmid (0.7 μg) and each sgRNA expression plasmid(0.3 μg) using Lipofectamine 2000 (Invitrogen) according to the manufacturer’s protocol. Cells were collected 5 days after transfection and the cell’s genomic DNA was prepared using NucleoSpin Tissue (MACHEREY-NAGEL & Co. KG).

GM14867 (*XPC* mutant fibroblasts) were purchased from Coriell Institute and maintained in Eagle's Minimum Essential Medium (EMEM) with 15% FBS and a penicillin/streptomycin mix. BJ-5ta cells (cat. no. CRL-4001, ATCC) were maintained in a 4:1 mixture of DMEM and Medium 199 with 10 μg/mL hygromycin B and 10% FBS. ARPE-19 cells (cat. no. CRL-2302, ATCC) were maintained in DMEM:F12 with 10% FBS and a penicillin/streptomycin mix. For plasmid-mediated expression of ABEs and sgRNAs, 6 × 10^5^ fibroblasts were co-transfected with 14 ug of ABE-encoding plasmid and 6 ug of sgRNA-expressing plasmid. Fibroblasts were transfected with the Amaxa P3 Primary Cell 4D-Nucleofector Kit using Program DS-137, according to the manufacturer’s protocol. A to G substitutions were analyzed 5 days after transfection.

### EGFP-PTC-knockin cell lines

2.5 × 10^5^ HeLa cells were transfected with Cas9-encoding plasmid (0.35 ug), AAVS1-sgRNA-encoding plasmid (0.15 ug), and EGFP-PTC encoding plasmid (0.5 ug) using the Neon™ transfection system (Invitrogen) with the following parameters: pulse voltage, 1,005; pulse width, 35 ms; pulse number, 2. Seven days after transfection, the culture medium was changed to 150ug/ml hygromycin B (Thermo Fisher Scientifics, cat. no. 10687010)-containing HeLa cell culture medium. Seven days after hygromycin B treatment, single cells were selected and cultured. Single cell-derived clones were analyzed and used for further experiments.

### Fluorescence-activated cell sorting

Five day after transfection, ABE-treated cells were trypsinized and resuspended in phosphate buffered saline. Single-cell suspensions were analyzed using a FACSCanto II (BD Biosciences) installed at Hanyang LINC+ Equipment Center (Seoul).

### Targeted deep sequencing

Genomic DNA segments that encompass the nuclease target sites were amplified using Phusion polymerase (New England Biolabs). Equal amounts of the PCR amplicons were subjected to paired-end read sequencing using Illumina MiSeq at Bio-Medical Science Co. (South Korea). Rare sequence reads that constituted less than 0.005% of the total reads were excluded. Off-targets were selected by Cas-OFFinder (http://www.rgenome.net/cas-offinder/)^23^ and base substitutions were analyzed by BE-Analyzer (http://www.rgenome.net/be-analyzer/)^39^. Primer sequences and List of off-targets are in Supplementary table 5 and Supplementary table 6.

### Treatment with ataluren and gentamicin

GM-14867 cells and the cells treated with xABE and ABEmax were maintained in EMEM with 15% FBS. When the confluency was 60-70%, the cells were treated with ataluren (10 μM; cat. no. S6003, Selleck) or gentamicin (1 mg/mL; cat. no. G1397, Sigma) for 48 h.

### Western blotting

Cell lysates were homogenized in 1X cell lysis buffer (cat. no. #9803, Cell Signaling Technology) and the supernatants were collected after centrifugation for 10 minutes at 14,000 *g*. An equal amount (35 μg) of the protein was separated by SDS-PAGE in 4–15% Mini-PROTEAN^®^ TGX™ Precast Protein Gels (cat. no. 4561084, Bio-Rad) and transferred to nitrocellulose membranes. The membranes were incubated with primary antibodies overnight at 4°C. The primary antibodies utilized in this study were as follows: anti-XPC antibody (cat. no. MA1-23328, Thermo), and anti-β-actin antibody (catalog no. A2668, Sigma). Then, the membranes were treated with the appropriate species-specific secondary antibodies (cat. no. sc-2357 and sc-516102, Santa Cruz) for 1 h at room temperature. After treatment of the membranes with reagents from the EZ-Western Lumi Pico Kit (cat. no. DG-WP100, DoGEN), the protein bands were visualized using the ImageQuant LAS4000 system with the accompanying software program (GE).

### Functional assessment

To assess the functional recovery of GM14867 cells, which carry a homozygous mutation in the *XPC* gene, after treatment with xABE, ABEmax, ataluren, or gentamicin, these cells, together with BJ-5ta WT cells, were exposed to ultraviolet irradiation at 254 nm at 1 J/m^2^/sec (cat. no. CL-1000, Analytik Jena) and left to grow for 72 h. Cells treated with ataluren or gentamicin underwent treatment for 48 h before ultraviolet exposure. Cell survival was evaluated with a water-soluble tetrazolium salt assay using an EZ-Cytox kit (cat. no. EZ-1000, DoGEN).

### Statistics

All statistical analyses were performed using the GraphPad Prism 5 program (GraphPad) and results are indicated in the figure legends. The values of each mean and standard error of mean were visualized as horizontal lines and error bars, respectively, in graphs.

## Acknowledgments

This work was supported by National Research Foundation of Korea (NRF) Grants (No. 2017R1A6A3A04004741 to D.H.J.; No. 2015M3A9E6028949, 2017M3A9B4062654, and 2018M3D1A1058826 to Je.H.K.; No. 2018M3A9H3022412 to S.B.), and by a grant from Korea Research Institute of Standards and Science (KRISS – 2018 – GP2018-0018) to Je.H.K. and by grants from Next Generation BioGreen 21 Program (PJ01319301), Korea Healthcare technology R&D Project (HI16C1012), Technology Innovation Program (No. 20000158) to S.B.

## Author contributions

Je.H.K. and S.B. conceived this project; C.L., D.H.J., J.Y., and S.P. performed the experiments; G.H.H. performed bioinformatics analyses; C.L., D.H.J., J.Y., and Ji.H.K. analyzed the data; J.S.K. gave critical comments; C.L., D.H.J., Je.H.K., and S.B. wrote the manuscript with the approval of all other authors.

## Data availability

Sequencing data has been uploaded to the Sequence Read Archive under Bioproject accession code PRJNA518883. All other data are available from the authors upon reasonable request.

## Additional information

Supplementary Information accompanies this papers at http://

## Competing interests

The authors declare no competing financial interests.

## REFERENCES

1. Alter J, et al. Systemic delivery of morpholino oligonucleotide restores dystrophin expression bodywide and improves dystrophic pathology. Nat Med 12, 175–177 (2006).

2. Aartsma-Rus A, van Ommen GJ. Antisense-mediated exon skipping: a versatile tool with therapeutic and research applications. RNA 13, 1609–1624 (2007).

3. Roy B, et al. Ataluren stimulates ribosomal selection of near-cognate tRNAs to promote nonsense suppression. Proc Natl Acad Sci U S A 113, 12508–12513 (2016).

4. Siddiqui N, Sonenberg N. Proposing a mechanism of action for ataluren. Proc Natl Acad Sci U S A 113, 12353–12355 (2016).

5. Kuschal C, DiGiovanna JJ, Khan SG, Gatti RA, Kraemer KH. Repair of UV photolesions in xeroderma pigmentosum group C cells induced by translational readthrough of premature termination codons. P Natl Acad Sci USA 110, 19483–19488 (2013).

6. Singh A, Ursic D, Davies J. Phenotypic suppression and misreading Saccharomyces cerevisiae. Nature 277, 146–148 (1979).

7. Pichavant C, et al. Current status of pharmaceutical and genetic therapeutic approaches to treat DMD. Mol Ther 19, 830–840 (2011).

8. Kim JS. Precision genome engineering through adenine and cytosine base editing. Nat Plants 4, 148–151 (2018).

9. Komor AC, Badran AH, Liu DR. Editing the Genome Without Double-Stranded DNA Breaks. ACS Chem Biol 13, 383–388 (2018).

10. Nami F, Basiri M, Satarian L, Curtiss C, Baharvand H, Verfaillie C. Strategies for In Vivo Genome Editing in Nondividing Cells. Trends Biotechnol 36, 770–786 (2018).

11. Komor AC, Kim YB, Packer MS, Zuris JA, Liu DR. Programmable editing of a target base in genomic DNA without double-stranded DNA cleavage. Nature 533, 420–424 (2016).

12. Nishida K, et al. Targeted nucleotide editing using hybrid prokaryotic and vertebrate adaptive immune systems. Science 353, aaf8729 (2016).

13. Gaudelli NM, et al. Programmable base editing of A*T to G*C in genomic DNA without DNA cleavage. Nature 551, 464–471 (2017).

14. Koblan LW, et al. Improving cytidine and adenine base editors by expression optimization and ancestral reconstruction. Nature biotechnol 36(9), 843–846 (2018).

15. Hu JH, et al. Evolved Cas9 variants with broad PAM compatibility and high DNA specificity. Nature 556, 57–63 (2018).

16. Nishimasu H, et al. Engineered CRISPR-Cas9 nuclease with expanded targeting space. Science 361, 1259–1262 (2018).

17. Li C, et al. Expanded base editing in rice and wheat using a Cas9-adenosine deaminase fusion. Genome Biol 19(1):59, (2018).

18. Ryu SM, et al. Adenine base editing in mouse embryos and an adult mouse model of Duchenne muscular dystrophy. Nat Biotechnol 36, 536–539 (2018).

19. DeKelver RC, et al. Functional genomics, proteomics, and regulatory DNA analysis in isogenic settings using zinc finger nuclease-driven transgenesis into a safe harbor locus in the human genome. Genome Res 20, 1133–1142 (2010).

20. Maresca M, Lin VG, Guo N, Yang Y. Obligate Ligation-Gated Recombination (ObLiGaRe): Custom-designed nuclease-mediated targeted integration through nonhomologous end joining. Genome Res 23, 539–546 (2013).

21. Sugasawa K, et al. Xeroderma Pigmentosum Group C Protein Complex Is the Initiator of Global Genome Nucleotide Excision Repair. Mol Cell 2, 223–232 (1998).

22. Dupuy A, et al. Targeted gene therapy of xeroderma pigmentosum cells using meganuclease and TALEN. PloS one 8, e78678 (2013).

23. Bae S, Park J, Kim JS. Cas-OFFinder: a fast and versatile algorithm that searches for potential off-target sites of Cas9 RNA-guided endonucleases. Bioinformatics 30, 1473–1475 (2014).

24. Liu Z, et al. Efficient generation of mouse models of human diseases via ABE- and BE-mediated base editing. Nat Commun 9, (2018).

25. Liang P, et al. Genome-wide profiling of adenine base editor specificity by EndoV-seq. Nat Commun 10, 67 (2019).

26. Lee HK, et al. Targeting fidelity of adenine and cytosine base editors in mouse embryos. Nat Commun 9, (2018).

27. Kuscu C, et al. CRISPR-STOP: gene silencing through base-editing-induced nonsense mutations. Nat Methods 14, 710–712 (2017).

28. Billon P, et al. CRISPR-Mediated Base Editing Enables Efficient Disruption of Eukaryotic Genes through Induction of STOP Codons. Mol Cell 67, 1068–1079 (2017).

29. Warrick E, et al. Preclinical Corrective Gene Transfer in Xeroderma Pigmentosum Human Skin Stem Cells. Mol Ther 20, 798–807 (2012).

30. Hacein-Bey-Abina S, et al. LMO2-associated clonal T cell proliferation in two patients after gene therapy for SCID-X1. Science 302, 415–419 (2003).

31. Roos WP, Kaina B. DNA damage-induced cell death by apoptosis. Trends Mol Med 12, 440–450 (2006).

32. Hess GT, Tycko J, Yao D, Bassik MC. Methods and Applications of CRISPR-Mediated Base Editing in Eukaryotic Genomes. Mol Cell 68, 26–43 (2017).

33. Nelson CE, et al. In vivo genome editing improves muscle function in a mouse model of Duchenne muscular dystrophy. Science 351, 403–407 (2016).

34. Benchaouir R, et al. Restoration of human dystrophin following transplantation of exon-skipping-engineered DMD patient stem cells into dystrophic mice. Cell Stem Cell 1, 646–657 (2007).

35. Kosicki M, Tomberg K, Bradley A. Repair of double-strand breaks induced by CRISPR-Cas9 leads to large deletions and complex rearrangements. Nat Biotechnol 36, 765–771 (2018).

36. Backliwal G, Hildinger M, Chenuet S, Wulhfard S, De Jesus M, Wurm FM. Rational vector design and multi-pathway modulation of HEK 293E cells yield recombinant antibody titers exceeding 1 g/l by transient transfection under serum-free conditions. Nucleic Acids Res 36, (2008).

37. Nguyen TA, et al. Functional Anatomy of the Human Microprocessor. Cell 161, 1374–1387 (2015).

38. Gibson DG, Young L, Chuang RY, Venter JC, Hutchison CA, 3rd, Smith HO. Enzymatic assembly of DNA molecules up to several hundred kilobases. Nat Methods 6, 343–345 (2009).

39. Hwang GH, et al. Web-based design and analysis tools for CRISPR base editing. Bmc Bioinformatics 19, (2018).

