## Supplementary Information for "CRISPR-pass: Gene rescue of nonsense mutations using adenine base editors"

**Content:**

**Supplementary Figure 1.** Coding or noncoding targeting depends on the *EGFP* sequence and the PTC position.

**Supplementary Figure 2.** Rescued EGFP expression after treatment with ABEs.

**Supplementary Figure 3.** FACS results

**Supplementary Figure 4.** CRISPR-pass for XPC patient-derived fibroblasts.

**Supplementary Figure 5.** Prolonged expression of the XPC protein after ABE treatment.

**Supplementary Figure 6.** Off-target analysis for CRISPR-pass targeting XPC.

**Supplementary Table 1.** FACS results.

**Supplementary Table 2.** NGS results.

**Supplementary Table 3.** A-to-G substitution rates (%) in potential ABE off-target sites.

**Supplementary Table 4.** List of oligomers encoding sgRNAs.

**Supplementary Table 5.** PCR primers used in this study.

**Supplementary Table 6.** List of off-target sites.

**Supplementary Figure 1. Coding or noncoding targeting depends on the *EGFP* sequence and the PTC position.** The sequence of the *EGFP* gene is shown. ABE target sequences are underlined. Depending on the target strand, codons for Lys53 or Asp217 were mutated such that they became STOP codons. The codons that are mutated are shown in blue and the PAM sequences are shown in red.

EGFP-PTCs-knockin DNA sequences

```
  1 atggtgagcaagggcgaggagctgttcaccggggtggtgcccatcctggtcgagctggacggcgacgtaaacggcca
 78 caagttcagcgtgtccggcgagggcgagggcgatgccacctacggcaagctgaccctgaagttcatctgcaccaccg
  coding strand target sgRNA (53rd Lysine -> STOP)
155 gcaagctgcccgtgccctggcccaccctcgtgaccaccctgacctacggcgtgcagtgcttcagccgctaccccgac
232 cacatgaagcagcacgacttcttcaagtccgccatgcccgaaggctacgtccaggagcgcaccatcttcttcaagga
309 cgacggcaactacaagacccgcgccgaggtgaagttcgagggcgacaccctggtgaaccgcatcgagctgaagggca
386 tcgacttcaaggaggacggcaacatcctggggcacaagctggagtacaactacaacagccacaacgtctatatcatg
463 gccgacaagcagaagaacggcatcaaggtgaacttcaagatccgccacaacatcgaggacggcagcgtgcagctcgc
540 cgaccactaccagcagaacacccccatcggcgacggccccgtgctgctgcccgacaaccactacctgagcacccagt
                    noncoding strand target sgRNA (217th Aspartic acid-> STOP)
617 ccgccctgagcaaagacccaacgagaagcgcgatcacatggtcctgctggagttcgtgaccgccgccgggatcact
694 ctcggcatggacgagctgtacaagtga
```

**Supplementary Figure 2. Rescued EGFP expression after treatment with ABEs.** Rescued EGFP expression in EGFP-PTC-KI cell lines in which the coding strand (a) or noncoding strand (b) is targeted for PTC bypass.

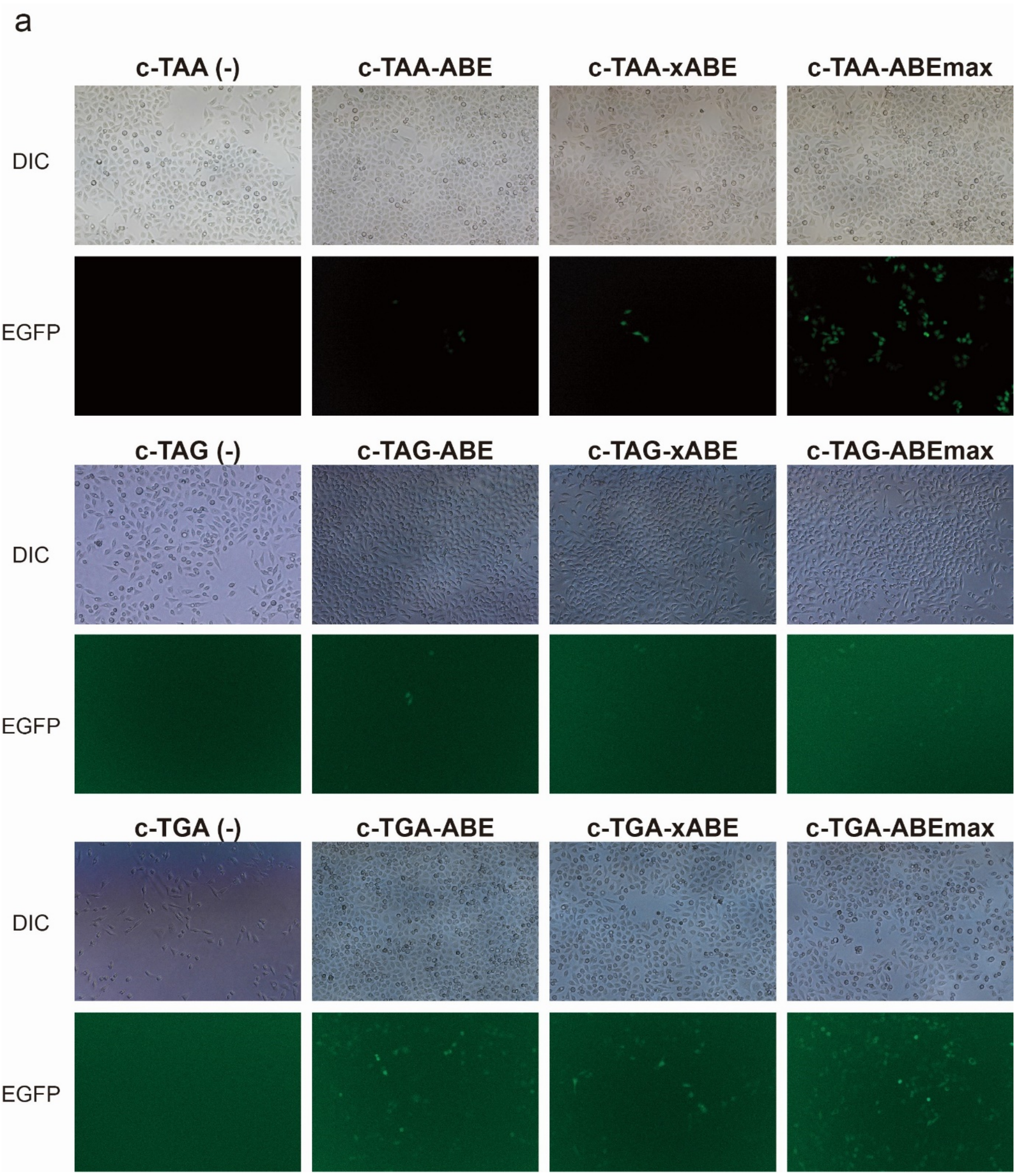

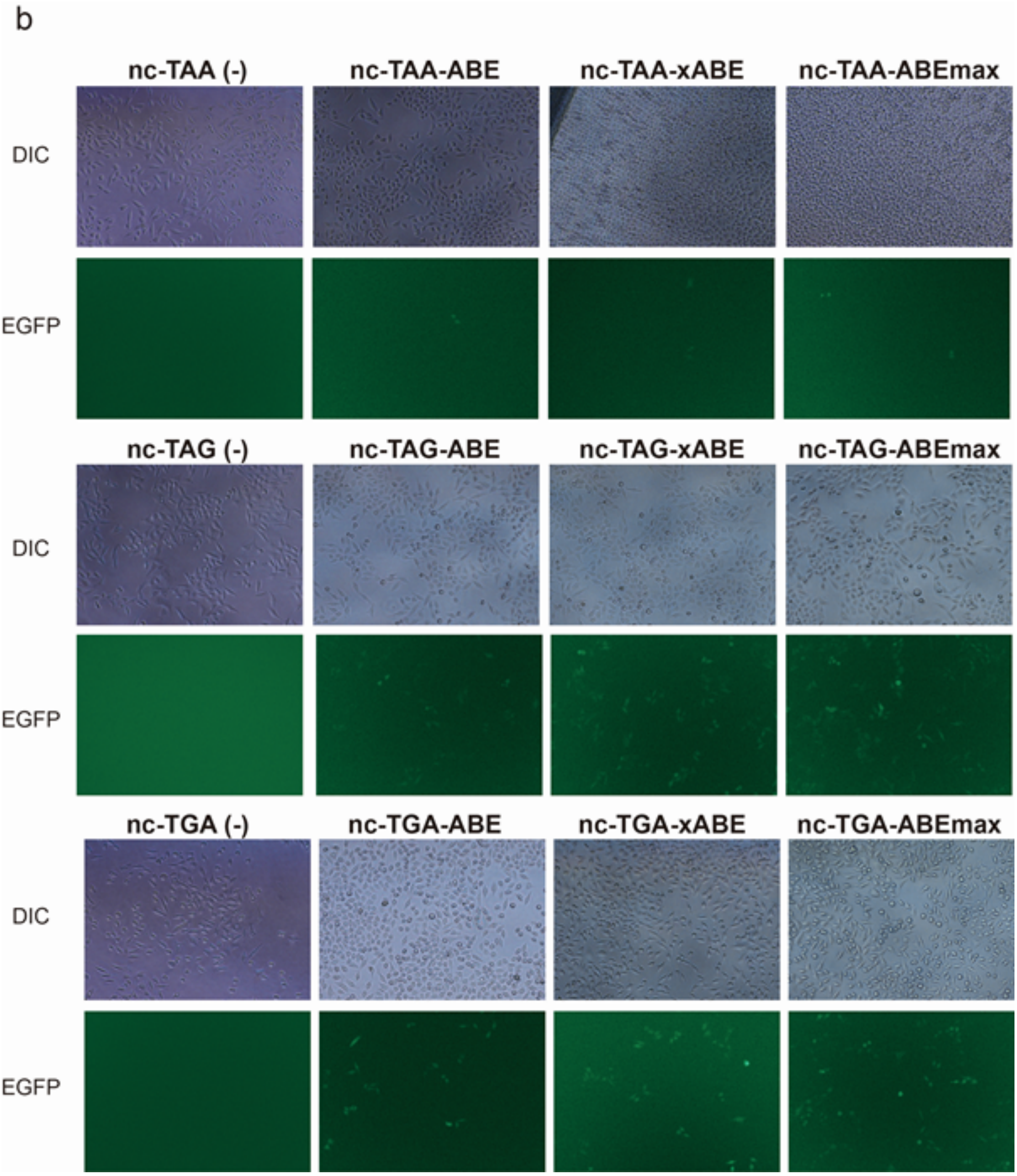
b
nc-TAA (-)
nc-TAA-ABE
nc-TAA-xABE
nc-TAA-ABEmax
DIC
EGFP
nc-TAG (-)
nc-TAG-ABE
nc-TAG-xABE
nc-TAG-ABEmax
DIC
EGFP
nc-TGA (-)
nc-TGA-ABE
nc-TGA-xABE
nc-TGA-ABEmax
DIC
EGFP

**Supplementary Figure 3. FACS results.**

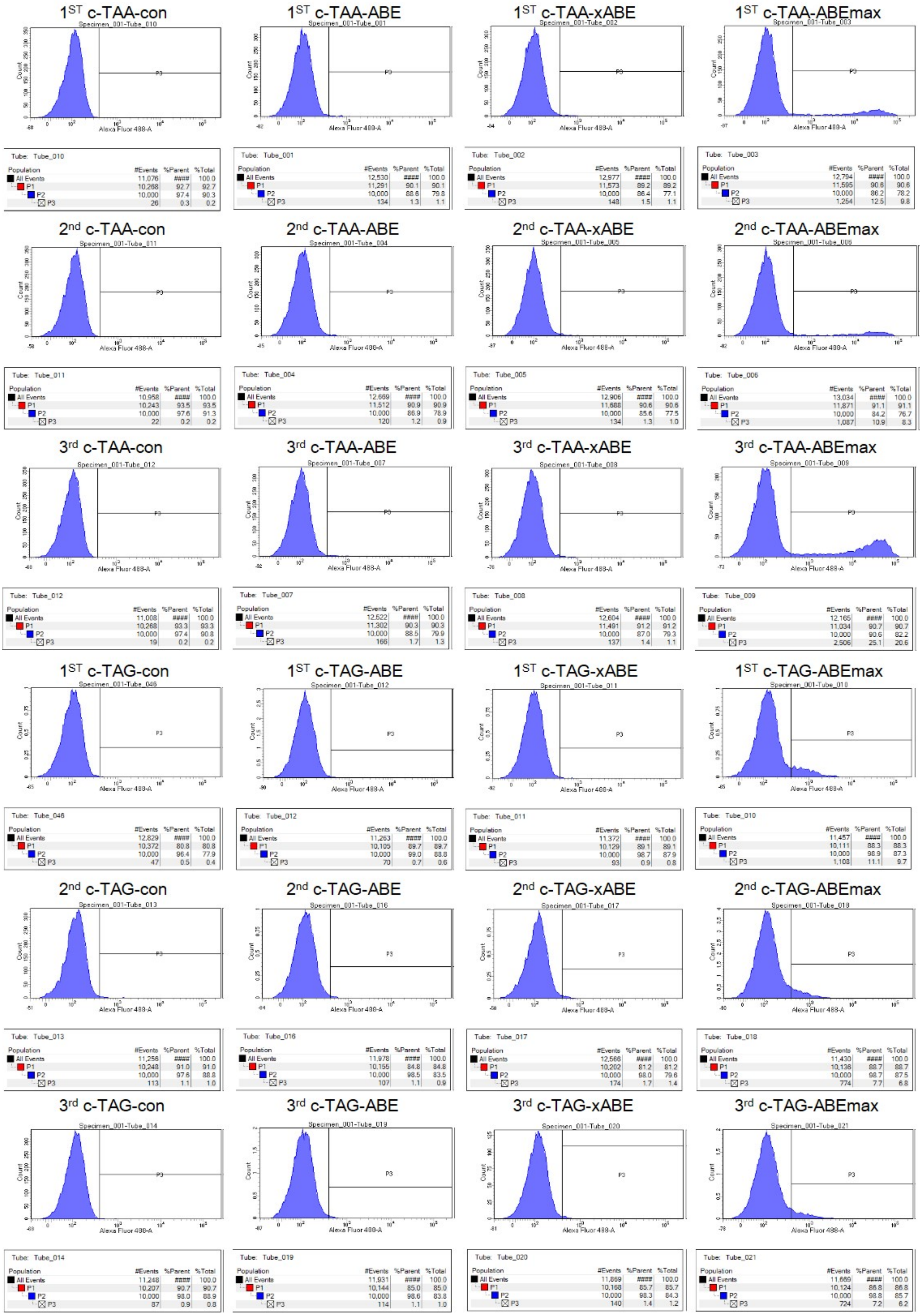

1$^{ST}$ c-TGA-con

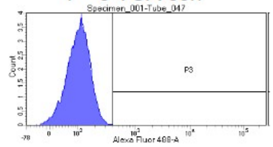


Tube: Tube_047

| Population | #Events | %Parent | %Total |
|---|---|---|---|
| All Events | 12,077 | #### | 100.0 |
| P1 | 10,212 | 84.6 | 84.6 |
| P2 | 10,000 | 97.9 | 82.8 |
| P3 | 43 | 0.4 | 0.4 |

1$^{ST}$ c-TGA-ABE

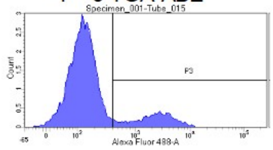


Tube: Tube_015

| Population | #Events | %Parent | %Total |
|---|---|---|---|
| All Events | 11,540 | #### | 100.0 |
| P1 | 10,131 | 87.8 | 87.8 |
| P2 | 10,000 | 98.7 | 86.7 |
| P3 | 1,784 | 17.8 | 15.5 |

1$^{ST}$ c-TGA-xABE

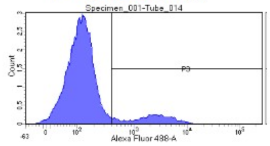


Tube: Tube_014

| Population | #Events | %Parent | %Total |
|---|---|---|---|
| All Events | 11,610 | #### | 100.0 |
| P1 | 10,176 | 87.6 | 87.6 |
| P2 | 10,032 | 98.6 | 86.4 |
| P3 | 1,315 | 13.1 | 11.3 |

1$^{ST}$ c-TGA-ABEmax

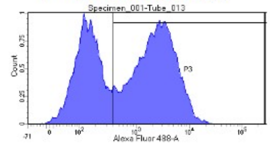


Tube: Tube_013

| Population | #Events | %Parent | %Total |
|---|---|---|---|
| All Events | 11,766 | #### | 100.0 |
| P1 | 10,153 | 86.3 | 86.3 |
| P2 | 10,000 | 98.5 | 85.0 |
| P3 | 5,962 | 59.6 | 50.7 |

2$^{nd}$ c-TGA-con

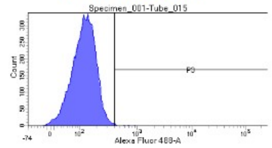


Tube: Tube_015

| Population | #Events | %Parent | %Total |
|---|---|---|---|
| All Events | 11,156 | #### | 100.0 |
| P1 | 10,185 | 91.3 | 91.3 |
| P2 | 10,000 | 98.2 | 89.6 |
| P3 | 78 | 0.8 | 0.7 |

2$^{nd}$ c-TGA-ABE

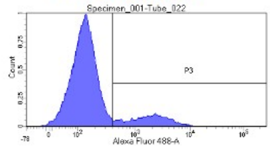


Tube: Tube_022

| Population | #Events | %Parent | %Total |
|---|---|---|---|
| All Events | 11,931 | #### | 100.0 |
| P1 | 10,169 | 85.2 | 85.2 |
| P2 | 10,000 | 98.3 | 83.8 |
| P3 | 1,622 | 16.2 | 13.6 |

2$^{nd}$ c-TGA-xABE

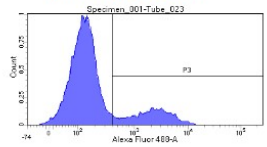


Tube: Tube_023

| Population | #Events | %Parent | %Total |
|---|---|---|---|
| All Events | 11,658 | #### | 100.0 |
| P1 | 10,150 | 87.1 | 87.1 |
| P2 | 10,000 | 98.5 | 85.8 |
| P3 | 2,035 | 20.3 | 17.5 |

2$^{nd}$ c-TGA-ABEmax

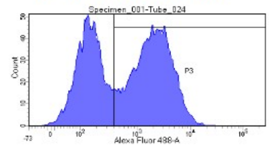


Tube: Tube_024

| Population | #Events | %Parent | %Total |
|---|---|---|---|
| All Events | 11,986 | #### | 100.0 |
| P1 | 10,242 | 85.4 | 85.4 |
| P2 | 10,000 | 97.6 | 83.4 |
| P3 | 5,771 | 57.7 | 48.1 |

3$^{rd}$ c-TGA-con

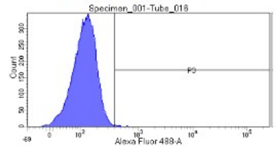


Tube: Tube_016

| Population | #Events | %Parent | %Total |
|---|---|---|---|
| All Events | 11,163 | #### | 100.0 |
| P1 | 10,205 | 91.4 | 91.4 |
| P2 | 10,000 | 98.0 | 89.6 |
| P3 | 81 | 0.8 | 0.7 |

3$^{rd}$ c-TGA-ABE

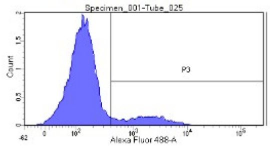


Tube: Tube_025

| Population | #Events | %Parent | %Total |
|---|---|---|---|
| All Events | 11,772 | #### | 100.0 |
| P1 | 10,158 | 86.3 | 86.3 |
| P2 | 10,000 | 98.4 | 84.9 |
| P3 | 1,418 | 14.2 | 12.0 |

3$^{rd}$ c-TGA-xABE

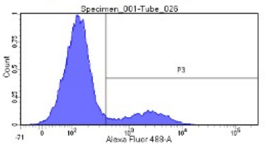


Tube: Tube_026

| Population | #Events | %Parent | %Total |
|---|---|---|---|
| All Events | 11,737 | #### | 100.0 |
| P1 | 10,161 | 86.6 | 86.6 |
| P2 | 10,000 | 98.4 | 85.2 |
| P3 | 1,823 | 18.2 | 15.5 |

3$^{rd}$ c-TGA-ABEmax

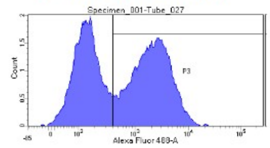


Tube: Tube_027

| Population | #Events | %Parent | %Total |
|---|---|---|---|
| All Events | 11,939 | #### | 100.0 |
| P1 | 10,209 | 85.5 | 85.5 |
| P2 | 10,000 | 98.0 | 83.8 |
| P3 | 5,502 | 55.0 | 46.1 |

1$^{ST}$ nc-TAA-con

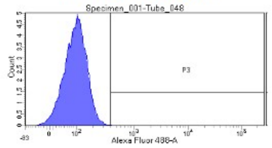


Tube: Tube_048

| Population | #Events | %Parent | %Total |
|---|---|---|---|
| All Events | 12,301 | #### | 100.0 |
| P1 | 10,486 | 85.2 | 85.2 |
| P2 | 10,000 | 95.4 | 81.3 |
| P3 | 34 | 0.3 | 0.3 |

1$^{ST}$ nc-TAA-ABE

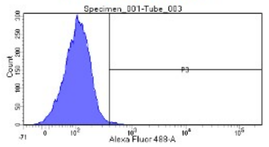


Tube: Tube_003

| Population | #Events | %Parent | %Total |
|---|---|---|---|
| All Events | 11,054 | #### | 100.0 |
| P1 | 9,667 | 87.5 | 87.5 |
| P2 | 9,528 | 98.6 | 86.2 |
| P3 | 134 | 1.4 | 1.2 |

1$^{ST}$ nc-TAA-xABE

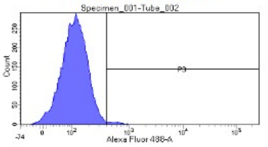


Tube: Tube_002

| Population | #Events | %Parent | %Total |
|---|---|---|---|
| All Events | 10,986 | #### | 100.0 |
| P1 | 9,689 | 88.2 | 88.2 |
| P2 | 9,529 | 98.3 | 86.7 |
| P3 | 180 | 1.9 | 1.6 |

1$^{ST}$ nc-TAA-ABEmax

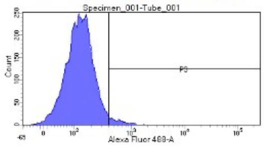


Tube: Tube_001

| Population | #Events | %Parent | %Total |
|---|---|---|---|
| All Events | 11,441 | #### | 100.0 |
| P1 | 9,054 | 79.1 | 79.1 |
| P2 | 8,873 | 98.0 | 77.6 |
| P3 | 307 | 3.5 | 2.7 |

2$^{nd}$ nc-TAA-con

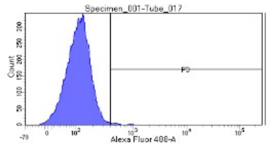


Tube: Tube_017

| Population | #Events | %Parent | %Total |
|---|---|---|---|
| All Events | 12,274 | #### | 100.0 |
| P1 | 10,554 | 86.0 | 86.0 |
| P2 | 10,000 | 94.8 | 81.5 |
| P3 | 140 | 1.4 | 1.1 |

2$^{nd}$ nc-TAA-ABE

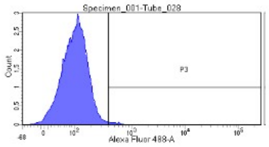


Tube: Tube_028

| Population | #Events | %Parent | %Total |
|---|---|---|---|
| All Events | 12,127 | #### | 100.0 |
| P1 | 10,296 | 84.9 | 84.9 |
| P2 | 10,000 | 97.1 | 82.5 |
| P3 | 133 | 1.3 | 1.1 |

2$^{nd}$ nc-TAA-xABE

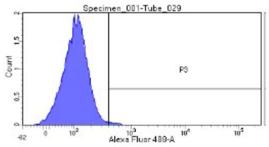


Tube: Tube_029

| Population | #Events | %Parent | %Total |
|---|---|---|---|
| All Events | 12,534 | #### | 100.0 |
| P1 | 10,415 | 83.1 | 83.1 |
| P2 | 10,000 | 96.0 | 79.8 |
| P3 | 129 | 1.3 | 1.0 |

2$^{nd}$ nc-TAA-ABEmax

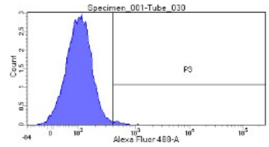


Tube: Tube_030

| Population | #Events | %Parent | %Total |
|---|---|---|---|
| All Events | 12,601 | #### | 100.0 |
| P1 | 10,469 | 83.1 | 83.1 |
| P2 | 10,000 | 95.5 | 79.4 |
| P3 | 180 | 1.8 | 1.4 |

3$^{rd}$ nc-TAA-con

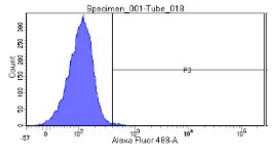


Tube: Tube_018

| Population | #Events | %Parent | %Total |
|---|---|---|---|
| All Events | 12,102 | #### | 100.0 |
| P1 | 10,466 | 86.5 | 86.5 |
| P2 | 10,000 | 95.5 | 82.6 |
| P3 | 136 | 1.4 | 1.1 |

3$^{rd}$ nc-TAA-ABE

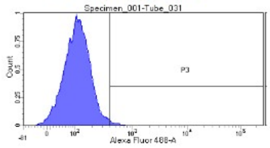


Tube: Tube_031

| Population | #Events | %Parent | %Total |
|---|---|---|---|
| All Events | 12,675 | #### | 100.0 |
| P1 | 10,183 | 80.3 | 80.3 |
| P2 | 10,000 | 98.2 | 78.9 |
| P3 | 143 | 1.4 | 1.1 |

3$^{rd}$ nc-TAA-xABE

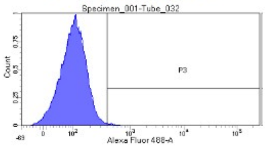


Tube: Tube_032

| Population | #Events | %Parent | %Total |
|---|---|---|---|
| All Events | 11,626 | #### | 100.0 |
| P1 | 10,120 | 87.0 | 87.0 |
| P2 | 10,000 | 98.8 | 86.0 |
| P3 | 98 | 1.0 | 0.8 |

3$^{rd}$ nc-TAA-ABEmax

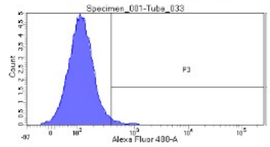


Tube: Tube_033

| Population | #Events | %Parent | %Total |
|---|---|---|---|
| All Events | 13,026 | #### | 100.0 |
| P1 | 10,249 | 78.7 | 78.7 |
| P2 | 10,000 | 97.6 | 76.8 |
| P3 | 257 | 2.6 | 2.0 |

### $1^{ST}$ nc-TAG-con

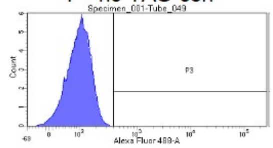


Tube: Tube_049

| Population | #Events | %Parent | %Total |
|---|---|---|---|
| All Events | 11,696 | #### | 100.0 |
| P1 | 10,203 | 87.2 | 87.2 |
| P2 | 10,000 | 98.0 | 85.5 |
| P3 | 23 | 0.2 | 0.2 |

### $1^{ST}$ nc-TAG-ABE

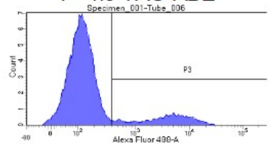


Tube: Tube_006

| Population | #Events | %Parent | %Total |
|---|---|---|---|
| All Events | 11,085 | #### | 100.0 |
| P1 | 10,117 | 91.3 | 91.3 |
| P2 | 10,000 | 98.8 | 90.2 |
| P3 | 1,707 | 17.1 | 15.4 |

### $1^{ST}$ nc-TAG-xABE

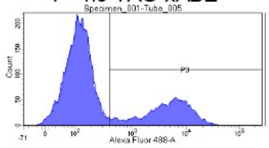


Tube: Tube_005

| Population | #Events | %Parent | %Total |
|---|---|---|---|
| All Events | 10,983 | #### | 100.0 |
| P1 | 10,109 | 92.0 | 92.0 |
| P2 | 10,000 | 98.9 | 91.0 |
| P3 | 3,036 | 30.4 | 27.6 |

### $1^{ST}$ nc-TAG-ABEmax

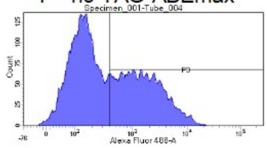


Tube: Tube_004

| Population | #Events | %Parent | %Total |
|---|---|---|---|
| All Events | 11,020 | #### | 100.0 |
| P1 | 9,627 | 87.4 | 87.4 |
| P2 | 9,438 | 98.0 | 85.6 |
| P3 | 4,210 | 44.6 | 38.2 |

### $2^{nd}$ nc-TAG-con

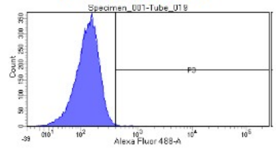


Tube: Tube_019

| Population | #Events | %Parent | %Total |
|---|---|---|---|
| All Events | 10,769 | #### | 100.0 |
| P1 | 10,116 | 93.9 | 93.9 |
| P2 | 10,000 | 98.9 | 92.9 |
| P3 | 92 | 0.9 | 0.9 |

### $2^{nd}$ nc-TAG-ABE

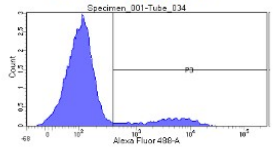


Tube: Tube_034

| Population | #Events | %Parent | %Total |
|---|---|---|---|
| All Events | 11,324 | #### | 100.0 |
| P1 | 10,196 | 90.0 | 90.0 |
| P2 | 10,000 | 98.1 | 88.3 |
| P3 | 1,368 | 13.7 | 12.1 |

### $2^{nd}$ nc-TAG-xABE

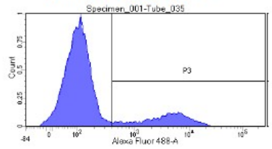


Tube: Tube_035

| Population | #Events | %Parent | %Total |
|---|---|---|---|
| All Events | 11,260 | #### | 100.0 |
| P1 | 10,179 | 90.4 | 90.4 |
| P2 | 10,000 | 98.2 | 88.8 |
| P3 | 1,978 | 19.8 | 17.6 |

### $2^{nd}$ nc-TAG-ABEmax

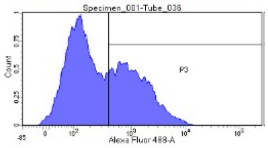


Tube: Tube_036

| Population | #Events | %Parent | %Total |
|---|---|---|---|
| All Events | 11,293 | #### | 100.0 |
| P1 | 10,172 | 90.1 | 90.1 |
| P2 | 10,000 | 98.3 | 88.6 |
| P3 | 4,160 | 41.6 | 36.8 |

### $3^{rd}$ nc-TAG-con

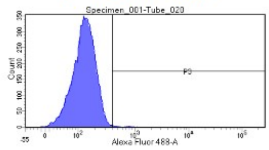


Tube: Tube_020

| Population | #Events | %Parent | %Total |
|---|---|---|---|
| All Events | 10,790 | #### | 100.0 |
| P1 | 10,132 | 93.9 | 93.9 |
| P2 | 10,000 | 98.7 | 92.7 |
| P3 | 131 | 1.3 | 1.2 |

### $3^{rd}$ nc-TAG-ABE

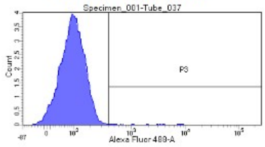


Tube: Tube_037

| Population | #Events | %Parent | %Total |
|---|---|---|---|
| All Events | 11,191 | #### | 100.0 |
| P1 | 10,194 | 91.1 | 91.1 |
| P2 | 10,000 | 98.1 | 89.4 |
| P3 | 279 | 2.8 | 2.5 |

### $3^{rd}$ nc-TAG-xABE

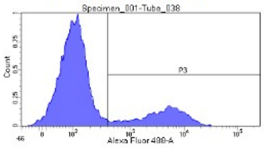


Tube: Tube_038

| Population | #Events | %Parent | %Total |
|---|---|---|---|
| All Events | 11,321 | #### | 100.0 |
| P1 | 10,207 | 90.2 | 90.2 |
| P2 | 10,000 | 98.0 | 88.3 |
| P3 | 2,294 | 22.9 | 20.3 |

### $3^{rd}$ nc-TAG-ABEmax

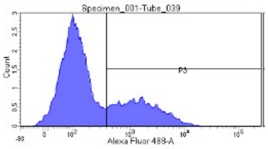


Tube: Tube_039

| Population | #Events | %Parent | %Total |
|---|---|---|---|
| All Events | 11,434 | #### | 100.0 |
| P1 | 10,147 | 88.7 | 88.7 |
| P2 | 10,000 | 98.6 | 87.5 |
| P3 | 3,137 | 31.4 | 27.4 |

### $1^{ST}$ nc-TGA-con

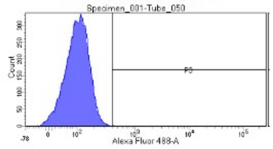


Tube: Tube_050

| Population | #Events | %Parent | %Total |
|---|---|---|---|
| All Events | 12,813 | #### | 100.0 |
| P1 | 10,354 | 80.8 | 80.8 |
| P2 | 10,000 | 96.6 | 78.0 |
| P3 | 36 | 0.4 | 0.3 |

### $1^{ST}$ nc-TGA-ABE

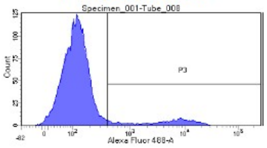


Tube: Tube_008

| Population | #Events | %Parent | %Total |
|---|---|---|---|
| All Events | 11,699 | #### | 100.0 |
| P1 | 10,179 | 87.0 | 87.0 |
| P2 | 10,000 | 98.2 | 85.5 |
| P3 | 1,200 | 12.0 | 10.3 |

### $1^{ST}$ nc-TGA-xABE

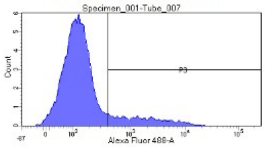


Tube: Tube_007

| Population | #Events | %Parent | %Total |
|---|---|---|---|
| All Events | 12,326 | #### | 100.0 |
| P1 | 10,196 | 82.7 | 82.7 |
| P2 | 10,000 | 98.1 | 81.1 |
| P3 | 1,573 | 15.7 | 12.8 |

### $1^{ST}$ nc-TGA-ABEmax

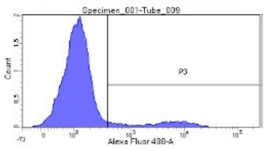


Tube: Tube_009

| Population | #Events | %Parent | %Total |
|---|---|---|---|
| All Events | 11,782 | #### | 100.0 |
| P1 | 10,178 | 86.4 | 86.4 |
| P2 | 10,000 | 98.3 | 84.9 |
| P3 | 1,099 | 11.0 | 9.3 |

### $2^{nd}$ nc-TGA-con

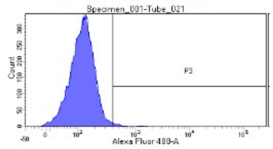


Tube: Tube_021

| Population | #Events | %Parent | %Total |
|---|---|---|---|
| All Events | 11,531 | #### | 100.0 |
| P1 | 10,237 | 88.8 | 88.8 |
| P2 | 10,000 | 97.7 | 86.7 |
| P3 | 206 | 2.1 | 1.8 |

### $2^{nd}$ nc-TGA-ABE

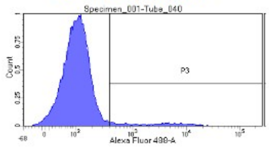


Tube: Tube_040

| Population | #Events | %Parent | %Total |
|---|---|---|---|
| All Events | 12,256 | #### | 100.0 |
| P1 | 10,278 | 83.9 | 83.9 |
| P2 | 10,000 | 97.3 | 81.6 |
| P3 | 689 | 6.9 | 5.6 |

### $2^{nd}$ nc-TGA-xABE

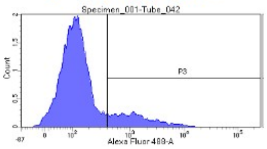


Tube: Tube_042

| Population | #Events | %Parent | %Total |
|---|---|---|---|
| All Events | 12,209 | #### | 100.0 |
| P1 | 10,260 | 84.0 | 84.0 |
| P2 | 10,000 | 97.5 | 81.9 |
| P3 | 1,726 | 17.3 | 14.1 |

### $2^{nd}$ nc-TGA-ABEmax

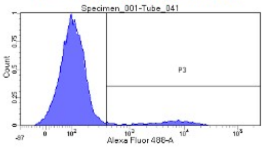


Tube: Tube_041

| Population | #Events | %Parent | %Total |
|---|---|---|---|
| All Events | 12,254 | #### | 100.0 |
| P1 | 10,274 | 83.8 | 83.8 |
| P2 | 10,000 | 97.3 | 81.6 |
| P3 | 950 | 9.5 | 7.8 |

### $3^{rd}$ nc-TGA-con

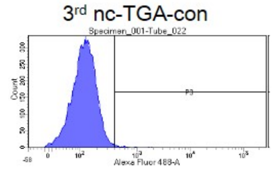


Tube: Tube_022

| Population | #Events | %Parent | %Total |
|---|---|---|---|
| All Events | 11,486 | #### | 100.0 |
| P1 | 10,221 | 89.0 | 89.0 |
| P2 | 10,000 | 97.8 | 87.1 |
| P3 | 161 | 1.6 | 1.4 |

### $3^{rd}$ nc-TGA-ABE

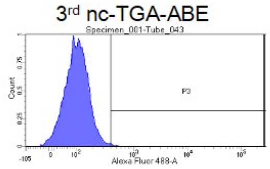


Tube: Tube_043

| Population | #Events | %Parent | %Total |
|---|---|---|---|
| All Events | 11,838 | #### | 100.0 |
| P1 | 10,216 | 86.3 | 86.3 |
| P2 | 10,000 | 97.9 | 84.5 |
| P3 | 182 | 1.8 | 1.5 |

### $3^{rd}$ nc-TGA-xABE

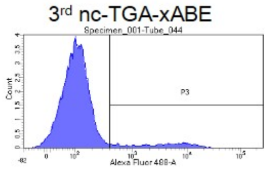


Tube: Tube_044

| Population | #Events | %Parent | %Total |
|---|---|---|---|
| All Events | 12,333 | #### | 100.0 |
| P1 | 10,287 | 83.4 | 83.4 |
| P2 | 10,000 | 97.2 | 81.1 |
| P3 | 1,015 | 10.2 | 8.2 |

### $3^{rd}$ nc-TGA-ABEmax

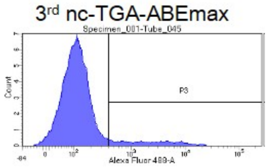


Tube: Tube_045

| Population | #Events | %Parent | %Total |
|---|---|---|---|
| All Events | 12,054 | #### | 100.0 |
| P1 | 10,216 | 84.8 | 84.8 |
| P2 | 10,000 | 97.9 | 83.0 |
| P3 | 1,129 | 11.3 | 9.4 |

**Supplementary Figure 4. CRISPR-pass for XPC patient-derived fibroblasts.** Next generation sequencing (NGS) results from (a) untreated GM14867 fibroblasts, (b) ABE-treated GM143867 fibroblasts, and (c) xABE-treated GM143867 fibroblasts. (d) Image of complete SDS-PAGE gel that is shown in part in Figure 3c.

a

*XPC*_wt

| | Trp | | Val | | | STOP | | | Asp | | | Val | | | Thr | | | Gln | | | Arg | | |
|---|---|---|---|---|---|---|---|---|---|---|---|---|---|---|---|---|---|---|---|---|---|---|---|
| | G | G | G | T | C | T | G | A | G | A | T | G | T | C | A | C | A | C | A | G | A | G | G |
| A | 0.1 | 0.1 | 0.1 | 0.0 | 0.0 | 0.0 | 0.0 | 99.7 | 0.0 | 99.7 | 0.0 | 0.1 | 0.0 | 0.0 | 99.7 | 0.0 | 99.8 | 0.0 | 99.8 | 0.0 | 99.7 | 0.1 | 0.1 |
| T | 0.0 | 0.0 | 0.0 | 99.6 | 0.0 | 99.8 | 0.0 | 0.0 | 0.0 | 0.0 | 99.7 | 0.0 | 99.6 | 0.1 | 0.0 | 0.1 | 0.0 | 0.0 | 0.0 | 0.0 | 0.0 | 0.0 | 0.0 |
| G | 99.9 | 99.9 | 99.9 | 0.1 | 0.0 | 0.0 | 100 | 0.2 | 100 | 0.2 | 0.0 | 99.9 | 0.0 | 0.0 | 0.2 | 0.0 | 0.2 | 0.0 | 0.2 | 100 | 0.3 | 99.9 | 99.9 |
| C | 0.0 | 0.0 | 0.0 | 0.3 | 100 | 0.2 | 0.0 | 0.0 | 0.0 | 0.0 | 0.3 | 0.0 | 0.4 | 99.9 | 0.0 | 99.9 | 0.0 | 100 | 0.0 | 0.0 | 0.0 | 0.0 | 0.0 |

b

*XPC*_ABE

| | Trp | | Val | | | STOP | | | Asp | | | Val | | | Thr | | | Gln | | | Arg | | |
|---|---|---|---|---|---|---|---|---|---|---|---|---|---|---|---|---|---|---|---|---|---|---|---|
| | G | G | G | T | C | T | G | A | G | A | T | G | T | C | A | C | A | C | A | G | A | G | G |
| A | 0.1 | 0.1 | 0.1 | 0.0 | 0.0 | 0.0 | 0.1 | 96.5 | 0.1 | 99.3 | 0.0 | 0.0 | 0.0 | 0.0 | 99.6 | 0.0 | 99.7 | 0.0 | 99.8 | 0.0 | 99.7 | 0.1 | 0.1 |
| T | 0.0 | 0.0 | 0.0 | 99.5 | 0.4 | 99.8 | 0.0 | 0.1 | 0.0 | 0.0 | 99.7 | 0.0 | 99.5 | 0.1 | 0.0 | 0.0 | 0.0 | 0.1 | 0.0 | 0.0 | 0.0 | 0.0 | 0.0 |
| G | 99.9 | 99.9 | 99.9 | 0.0 | 0.0 | 0.0 | 99.9 | 3.4 | 99.9 | 0.7 | 0.0 | 99.9 | 0.0 | 0.0 | 0.3 | 0.0 | 0.3 | 0.0 | 0.2 | 99.9 | 0.3 | 99.9 | 99.9 |
| C | 0.0 | 0.0 | 0.0 | 0.4 | 99.6 | 0.2 | 0.0 | 0.0 | 0.0 | 0.0 | 0.3 | 0.0 | 0.4 | 99.9 | 0.0 | 99.9 | 0.0 | 99.9 | 0.0 | 0.0 | 0.0 | 0.0 | 0.0 |

c

*XPC*_xABE

| | Trp | | Val | | | STOP | | | Asp | | | Val | | | Thr | | | Gln | | | Arg | | |
|---|---|---|---|---|---|---|---|---|---|---|---|---|---|---|---|---|---|---|---|---|---|---|---|
| | G | G | G | T | C | T | G | A | G | A | T | G | T | C | A | C | A | C | A | G | A | G | G |
| A | 0.1 | 0.1 | 0.1 | 0.0 | 0.0 | 0.0 | 0.0 | 86.2 | 0.0 | 99.3 | 0.0 | 0.0 | 0.0 | 0.0 | 99.7 | 0.0 | 99.7 | 0.0 | 99.8 | 0.0 | 99.6 | 0.1 | 0.1 |
| T | 0.0 | 0.0 | 0.0 | 99.5 | 0.7 | 99.7 | 0.0 | 0.1 | 0.0 | 0.0 | 99.7 | 0.0 | 99.5 | 0.0 | 0.0 | 0.1 | 0.0 | 0.1 | 0.0 | 0.0 | 0.1 | 0.0 | 0.0 |
| G | 99.9 | 99.9 | 99.9 | 0.0 | 0.1 | 0.0 | 99.9 | 13.7 | 99.9 | 0.7 | 0.0 | 99.9 | 0.0 | 0.0 | 0.3 | 0.0 | 0.2 | 0.0 | 0.2 | 100 | 0.4 | 99.9 | 99.9 |
| C | 0.0 | 0.0 | 0.0 | 0.4 | 99.2 | 0.2 | 0.0 | 0.0 | 0.0 | 0.0 | 0.2 | 0.0 | 0.5 | 99.9 | 0.0 | 99.9 | 0.0 | 99.9 | 0.0 | 0.0 | 0.0 | 0.0 | 0.0 |

d

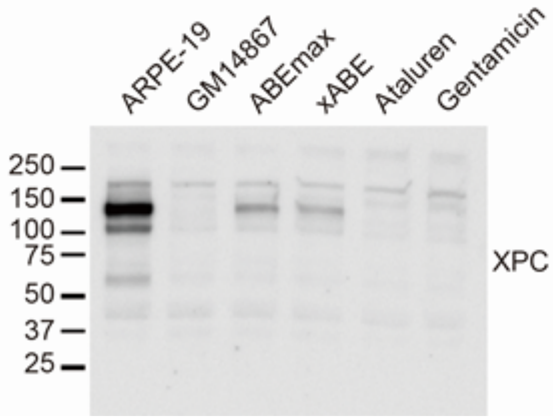

**Supplementary Figure 5. Prolonged expression of the XPC protein after ABE treatment.** Image of complete SDS-PAGE gel that is shown in part in Figure 3e.

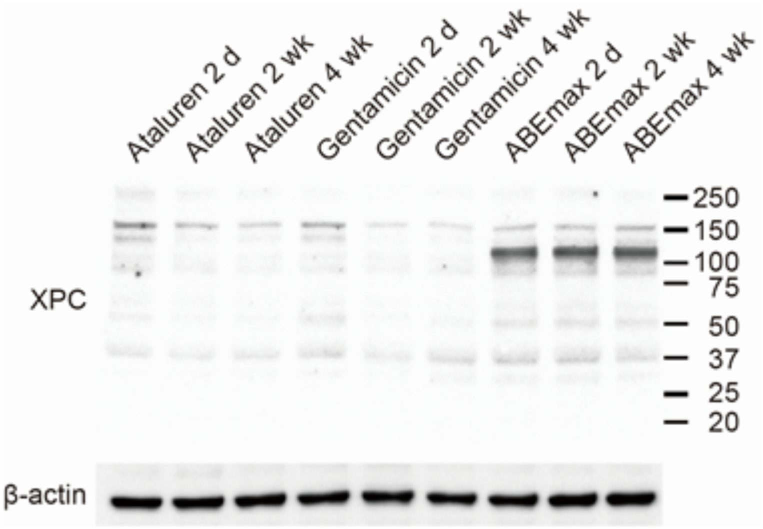

**Supplementary Figure 6. Off-target analysis for CRISPR-pass targeting XPC.** A to G substitution rates at off-target sites are displayed. The percentages of substitutions at each site are summarized in Supplementary Table 3. The red bars depict A to G substitution rates in untreated samples, whereas the blue bars depict A to G substitution rates in ABE-treated samples. Blue arrowhead indicates a target "A" which shows the A to G substitution rates (%).

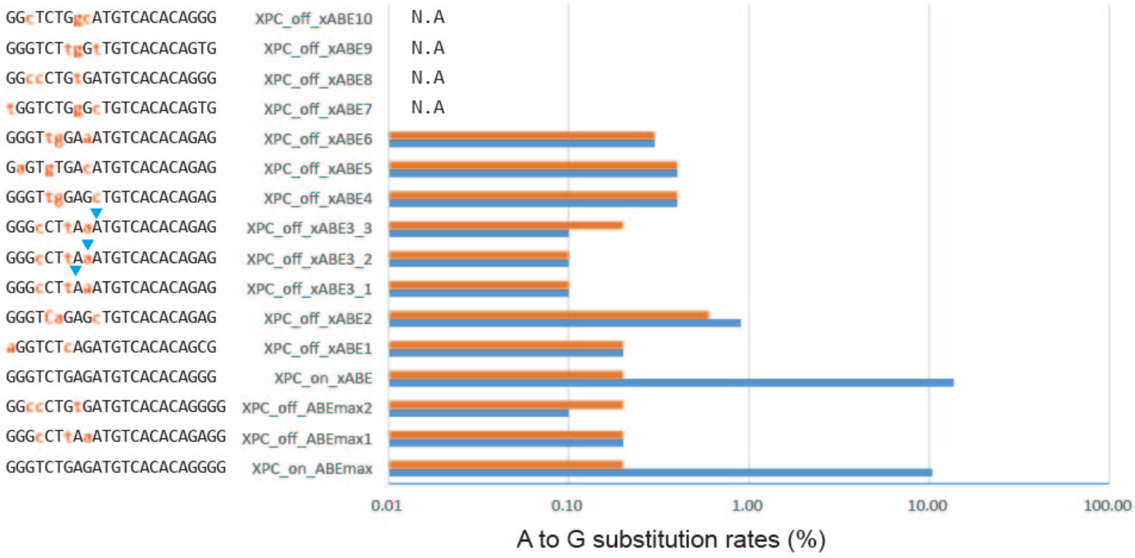

**Supplementary Table 1. FACS results.** The percentages of EGFP (+) cells in populations of ABE-treated EGFP-PTC-KI cells. Each experiment was repeated 3 times.

| | (-) | | | ABE | | | xABE | | | ABEmax | | |
|---|---|---|---|---|---|---|---|---|---|---|---|---|
| | 1st trial | 2nd trial | 3rd trial | 1st trial | 2nd trial | 3rd trial | 1st trial | 2nd trial | 3rd trial | 1st trial | 2nd trial | 3rd trial |
| c-TAA | 0.3% | 0.2% | 0.2% | 1.3% | 1.2% | 1.7% | 1.5% | 1.3% | 1.4% | 12.5% | 10.9% | 25.1% |
| c-TAG | 0.5% | 1.1% | 0.9% | 0.7% | 1.1% | 1.1% | 0.9% | 1.7% | 1.4% | 11.1% | 7.7% | 7.2% |
| c-TGA | 0.4% | 0.8% | 0.8% | 17.8% | 16.2% | 14.2% | 13.1% | 20.3% | 18.2% | 59.6% | 57.7% | 55.0% |
| nc-TAA | 0.3% | 1.4% | 1.4% | 1.4% | 1.3% | 1.4% | 1.9% | 1.3% | 1.0% | 3.5% | 1.8% | 2.6% |
| nc-TAG | 0.2% | 0.9% | 1.3% | 17.1% | 13.7% | 2.8% | 30.4% | 19.8% | 22.9% | 44.6% | 41.6% | 31.4% |
| nc-TGA | 0.4% | 2.1% | 1.6% | 11.0% | 6.9% | 1.8% | 12.0% | 9.5% | 10.2% | 15.7% | 17.3% | 11.3% |

**Supplementary Table 2. NGS results.** The percentages of A to G substitutions in populations of ABE-treated EGFP-PTC-KI cells. Each experiment was repeated 3 times.

| | (-) | | | ABE | | | xABE | | | ABEmax | | |
|---|---|---|---|---|---|---|---|---|---|---|---|---|
| | 1st trial | 2nd trial | 3rd trial | 1st trial | 2nd trial | 3rd trial | 1st trial | 2nd trial | 3rd trial | 1st trial | 2nd trial | 3rd trial |
| c-TAA | 0.0% | 0.1% | 0.0% | 0.2% | 0.2% | 0.0% | 0.0% | 0.0% | 0.9% | 11.2% | 9.7% | 43.5% |
| c-TAG | 0.5% | 0.6% | 0.4% | 5.3% | 6.5% | 2.7% | 9.1% | 9.5% | 4.0% | 51.2% | 52.2% | 54.1% |
| c-TGA | 0.3% | 0.3% | 0.0% | 6.2% | 5.9% | 5.4% | 5.1% | 4.4% | 2.4% | 37.5% | 34.0% | 41.3% |
| nc-TAA | 0.3% | 0.2% | 0.5% | 3.1% | 0.4% | 2.6% | 7.1% | 9.3% | 8.7% | 43.4% | 10.5% | 12.4% |
| nc-TAG | 0.3% | 0.1% | 0.2% | 12.7% | 1.8% | 14.3% | 16.3% | 22.2% | 28.7% | 41.9% | 33.1% | 44.8% |
| nc-TGA | 0.2% | 0.3% | 0.4% | 9.1% | 2.3% | 15.2% | 15.0% | 12.1% | 16.0% | 37.4% | 19.2% | 26.9% |

**Supplementary Table 3. A-to-G substitution rates (%) in potential ABE off-target sites.** N.A., not available; these sites are a Cas9 or xCas9 off-target site but do not contain an A targetable by ABEs. Blue colored letter means a target "A" which shows the A to G substitution rates (%)

| Name | Target sequences | ABE treated | wt |
|---|---|---|---|
| XPC_ABEmax_on | GGGTCTGAGATGTCACACAGNGG | 10.40% | 0.20% |
| XPC__ABEmax off_1 | GGGcCTtAaATGTCACACAGAGG | 0.20% | 0.20% |
| XPC__ABEmax off_2 | GGccCTGtGATGTCACACAGGGG | 0.10% | 0.20% |
| XPC_ xABE_on | GGGTCTGAGATGTCACACAGNG | 13.70% | 0.20% |
| XPC_ xABE_off_1 | aGGTCTcAGATGTCACACAGCG | 0.20% | 0.20% |
| XPC_ xABE_off_2 | GGGTCaGAGcTGTCACACAGAG | 0.90% | 0.60% |
| XPC_ xABE_off_3_1 | GGGcCTtAaATGTCACACAGAG | 0.10% | 0.10% |
| XPC_ xABE_off_3_2 | GGGcCTtAaATGTCACACAGAG | 0.10% | 0.10% |
| XPC_ xABE_off_3_3 | GGGcCTtAaATGTCACACAGAG | 0.10% | 0.20% |
| XPC_ xABE_off_4 | GGGTtgGAGcTGTCACACAGAG | 0.40% | 0.40% |
| XPC_ xABE_off_5 | GaGTgTGAcATGTCACACAGAG | 0.40% | 0.40% |
| XPC_ xABE_off_6 | GGGTtgGAaATGTCACACAGAG | 0.30% | 0.30% |
| XPC_ xABE_off_7 | tGGTCTGgGcTGTCACACAGTG | N.A | N.A |
| XPC_ xABE_off_8 | GGccCTGtGATGTCACACAGGG | N.A | N.A |
| XPC_ xABE_off_9 | GGGTCTtgGtTGTCACACAGTG | N.A | N.A |
| XPC_ xABE_off_10 | GGcTCTGgcATGTCACACAGGG | N.A | N.A |

**Supplementary Table 4. List of oligomers encoding sgRNAs.**

| Name | sequences |
|---|---|
| psg-nc-TAA-up | CACCGTCTCTTAGGGGTCTTTGCTC |
| psg-nc-TAG-up | CACCGTCTCCTAGGGGTCTTTGCTC |
| psg-nc-TGA-up | CACCGTCTCTCAGGGGTCTTTGCTC |
| psg-nc-TAA-bo | AAACGAGCAAAGACCCCTAAGAGAC |
| psg-nc-TAG-bo | AAACGAGCAAAGACCCCTAGGAGAC |
| psg-nc-TGA-bo | AAACGAGCAAAGACCCCTGAGAGAC |
| psg-c-TAG_1up | CACCGCCGGCTAGCTGCCCGTGCCC |
| psg-c-TAA_2up | CACCGCCGGCTAACTGCCCGTGCCC |
| psg-c-TGA_3up | CACCGCCGGCTGACTGCCCGTGCCC |
| psg-c-TAG_1bo | AAACGGGCACGGGCAGCTAGCCGGC |
| psg-c-TAA_2bo | AAACGGGCACGGGCAGTTAGCCGGC |
| psg-c-TGA_3bo | AAACGGGCACGGGCAGTCAGCCGGC |
| psg-XPC-up | CACCGGGGTCTGAGATGTCACACAG |
| psg-XPC-bo | AAACGACACACTGTAGAGACTGGGC |
| psg-AAVS1-up | CACCGTAAGCAAACCTTAGAGGTTC |
| psg-AAVS1-bo | AAACCTTGGAGATTCCAAACGAATC |

**Supplementary Table 5. PCR primers used in this study.**

| Name | Sequences |
|---|---|
| EGFP-1stF | gacatatccacgccctccta |
| EGFP-1stR | ctgacaattccgtggtgttg |
| EGFP_c_PTC_Deep_F | ACACTCTTTCCCTACACGAC GCTCTTCCGATCT acgtaaacggccacaagttc |
| EGFP_c_PTC_Deep_R | GTGACTGGAGTTCAGACGTGT GCTCTTCCGATCT tcgtccttgaagaagatggtg |
| EGFP_nc_PTC_Deep_F | ACACTCTTTCCCTACACGAC GCTCTTCCGATCT gaacggcatcaaggtgaact |
| EGFP_nc_PTC_Deep_R | GTGACTGGAGTTCAGACGTGT GCTCTTCCGATCT cttgtacagctcgtccatgc |
| inf_sacI_Cgo_add_F | GGTCTATATAAGCAGAGCTC TCGTCGACGAGCTCGTTTAGTG |
| inf_sacI_Cgo_add_R | CTCACCATGGCGGCGAGCTC GGTACCCTGGACACCTGTGG |
| inf_ccn_n2_TAA_2F | CTGAGCAAAGACCCCtgagagaagcgcgatcacatgg |
| inf_ccn_n2_TAA_1R | tcaGGGGTCTTTGCTCAGGGCG |
| inf_ccn_n2_TAG_2F | CTGAGCAAAGACCCCcaagagaagcgcgatcacatgg |
| inf_ccn_n2_TAG_1R | ttgGGGGTCTTTGCTCAGGGCG |
| inf_ccn_n2_TGA_2F | CTGAGCAAAGACCCCcgagagaagcgcgatcacatgg |
| inf_ccn_n2_TGA_1R | tcgGGGGTCTTTGCTCAGGGCG |
| inf_go_F1-1 | TGGGAGGTCTATATAAGCAGAGCTCATGGTGAGCAAGGGCGAGG |
| inf_go_R1-TAG | CCATGTGCTAGCGCTTCTCGTTGGGGTC |
| inf_go_R1-TAA | CCATGTGTTAGCGCTTCTCGTTGGGGTC |
| inf_go_R1-TGA | CCATGTGTCAGCGCTTCTCGTTGGGGTC |
| inf_go_F1-2TAG | CGAGAAGCGCTAGcacatggtcctgctggagtt |
| inf_go_F1-2TAA | CGAGAAGCGCTAAcacatggtcctgctggagtt |
| inf_go_F1-2TGA | CGAGAAGCGCTGAcacatggtcctgctggagtt |
| inf_go_R1-2 | TGAGATGTCTCTGTGCGGCTCACTTGTACAGCTCGTCCATGC |
| inf_go_R2-1TAG | CGGGCAGCTAGCCGGTGGTGCAGATGAAC |
| inf_go_R2-1TAA | CGGGCAGTTAGCCGGTGGTGCAGATGAAC |
| inf_go_R2-1TGA | CGGGCAGTCAGCCGGTGGTGCAGATGAAC |
| inf_go_F2-2TAG | CACCACCGGCTAGCTGCCCGTGCCCTGGCCC |
| inf_go_F2-2TAA | CACCACCGGCTAACTGCCCGTGCCCTGGCCC |
| inf_go_F2-2TGA | CACCACCGGCTGACTGCCCGTGCCCTGGCCC |

| Name | Sequences |
|---|---|
| XPC_1stF | ccaggagacaagcaggagaa |
| XPC_1stR | cgcggcagttcatctttcaa |
| XPC_deepF | ACACTCTTTCCCTACACGAC GCTCTTCCGATCT gtgagcaggaggaaaagtgg |
| XPC_deepR | GTGACTGGAGTTCAGACGTGT GCTCTTCCGATCT gtatggtctcaaggtctcggc |
| XPC_off_2nd_F1 | ACACTCTTTCCCTACACGAC GCTCTTCCGATCT CACATGCTCCTGGAAGGGAA |
| XPC_off_2nd_R1 | GTGACTGGAGTTCAGACGTGT GCTCTTCCGATCT AGGAGTGCCTACAGATGGGT |
| XPC_off_2nd_F2 | ACACTCTTTCCCTACACGAC GCTCTTCCGATCT TTCACAGGCTGGCATTGAGT |
| XPC_off_2nd_R2 | GTGACTGGAGTTCAGACGTGT GCTCTTCCGATCT TGCCCAGACAGAAGTTTGCT |
| XPC_off_2nd_F3_NGG_F1 | ACACTCTTTCCCTACACGAC GCTCTTCCGATCT TGGAAGTGTAAAGGGGTTGTCT |
| XPC_off_2nd_R3_NGG_R1 | GTGACTGGAGTTCAGACGTGT GCTCTTCCGATCT TCCATCTTTCACAGAGCTTCCA |
| XPC_off_2nd_F4 | ACACTCTTTCCCTACACGAC GCTCTTCCGATCT GCATTTCCAGGCACACAGTG |
| XPC_off_2nd_R4 | GTGACTGGAGTTCAGACGTGT GCTCTTCCGATCT CAGAGGATGCAAGGAAACACC |
| XPC_off_2nd_F5 | ACACTCTTTCCCTACACGAC GCTCTTCCGATCT TCCATTTAGCTCGGGATGGC |
| XPC_off_2nd_R5 | GTGACTGGAGTTCAGACGTGT GCTCTTCCGATCT TGCCTCATTGTTCATTAGTGTCT |
| XPC_off_2nd_F6 | ACACTCTTTCCCTACACGAC GCTCTTCCGATCT AGTCATAATATTTCAAGGCAGAAAAGA |
| XPC_off_2nd_R6 | GTGACTGGAGTTCAGACGTGT GCTCTTCCGATCT ACGCTCTTTTCAGACATTCTTGT |
| XPC_off_2nd_F7 | ACACTCTTTCCCTACACGAC GCTCTTCCGATCT TGGCAGCAAGAGAAAGGAGG |
| XPC_off_2nd_R7 | GTGACTGGAGTTCAGACGTGT GCTCTTCCGATCT GTGACCTTCCTCCTTCCGTG |
| XPC_off_2nd_F8_NGG_F1 | ACACTCTTTCCCTACACGAC GCTCTTCCGATCT GACCTGTACTATGGGCTGCC |
| XPC_off_2nd_R8_NGG R1 | GTGACTGGAGTTCAGACGTGT GCTCTTCCGATCT TCATCATCCCCTCCCTGTGT |
| XPC_off_2nd_F9 | ACACTCTTTCCCTACACGAC GCTCTTCCGATCT ACCTCCCTCCTGAAGAAGTGA |
| XPC_off_2nd_R9 | GTGACTGGAGTTCAGACGTGT GCTCTTCCGATCT TGGGCAGGACTGATATCCCT |
| XPC_off_2nd_F10 | ACACTCTTTCCCTACACGAC GCTCTTCCGATCT CCTCCTAAGGAACAACATGGTGT |
| XPC_off_2nd_R10 | GTGACTGGAGTTCAGACGTGT GCTCTTCCGATCT TGCAATTTCTTCTTTGTCCTGAGT |
| XPC_off_1st_F1 | TGCAAACCCCTTCTGTCTGT |
| XPC_off_1st_R1 | TGCAGTGAGCTGAGATTGGG |
| XPC_off_1st_F2 | AATGGGGGTACAGGCATTGG |
| XPC_off_1st_R2 | AGCTGGCTGCAGAAATTTGC |
| XPC_off_1st_F3_NGG_F1 | GAGGTTGCAGTGAGCCAAGA |
| XPC_off_1st_R3_NGG_R1 | GGAGGGAGAGAGGAGTGGAG |
| XPC_off_1st_F4 | GCCTTCTCAACAATCCCCCA |
| XPC_off_1st_R4 | CCACTGTTTTGTGCAGCCTC |
| XPC_off_1st_F5 | TGAGGCGTGGAAGTGTGTAC |
| XPC_off_1st_R5 | TCAGCTCACTGCAACCTCTG |
| XPC_off_1st_F6 | CTTACCAGCGGCTCTTGGAA |
| XPC_off_1st_R6 | CATCTGCTAAAGGGCTGGCT |
| XPC_off_1st_F7 | CCTCACAGCCAATCCCATGT |
| XPC_off_1st_R7 | AGGAGTGGCTCATCAAAGGC |
| XPC_off_1st_F8_NGG_F1 | ATGTGGACCCAGGCATTCTG |

| | |
|---|---|
| XPC_off_1st_R8_NGG_R1 | CAGAGGGAGCACCAAGGAAG |
| XPC_off_1st_F9 | GCAAGGGAGAAAGGAGGGTC |
| XPC_off_1st_R9 | CTCCTTCTTGTCGTGGGGAC |
| XPC_off_1st_F10 | TTCCAAACCCCCAGGAACTT |
| XPC_off_1st_R10 | TCAGCCATACCACACCAAGA |
